# Genomes of nitrogen-fixing eukaryotes reveal a non-canonical model of organellogenesis

**DOI:** 10.1101/2024.08.27.609708

**Authors:** Sarah Frail, Melissa Steele-Ogus, Jon Doenier, Solène L.Y. Moulin, Tom Braukmann, Shouling Xu, Ellen Yeh

**Affiliations:** Department of Biochemistry, Stanford School of Medicine, Stanford, CA 94305, USA; Department of Pathology, Stanford School of Medicine, Stanford, CA 94305, USA; Department of Plant Biology, Carnegie Institution, Stanford, CA 94305, USA; Department of Microbiology & Immunology, Stanford School of Medicine, Stanford, CA 94305, USA; Chan Zuckerberg Biohub – San Francisco, San Francisco, California 94158, USA

**Keywords:** Cyanobacteria, diatom, endosymbiosis, evolution, genome, genomics, horizontal gene transfer, nitrogen fixation, organelle, organellogenesis, symbiosis

## Abstract

Endosymbiotic gene transfer and import of host-encoded proteins are considered hallmarks of organelles necessary for stable integration of two cells. However, newer endosymbiotic models have challenged the origin and timing of such genetic integration during organellogenesis. Epithemia diatoms contain diazoplasts, obligate endosymbionts that are closely related to recently-described nitrogen-fixing organelles and share similar function as integral cell compartments. We report genomic analyses of two species which are highly divergent but share a common ancestor at the origin of the endosymbiosis. We found minimal evidence of genetic integration in E.clementina: nonfunctional diazoplast-to-nucleus DNA transfers and 6 host-encoded proteins of unknown function in the diazoplast proteome, far fewer than detected in other recently-acquired endosymbionts designated organelles. Epithemia diazoplasts are a valuable counterpoint to existing organellogenesis models, demonstrating that endosymbionts can function as integral compartments absent significant genetic integration. The minimal genetic integration makes diazoplasts valuable blueprints for bioengineering endosymbiotic compartments de novo.

## INTRODUCTION

Endosymbiotic organelles are uniquely eukaryotic innovations for the acquisition of complex cellular functions, including aerobic respiration, photosynthesis, and nitrogen fixation. Endosymbioses contributed to expansive eukaryotic diversity^1^. An important question in cell evolution and engineering is: how do intermittent, facultative endosymbioses evolve into permanent integral cell compartments, i.e. organelles? With the recognition of the bacterial origin of mitochondria and chloroplasts in the 1980s, Cavalier-Smith & Lee proposed that the key distinction between a transient endosymbiont and an organelle was that organelles do not synthesize all their own proteins^2,3^. Instead, in organelles, some genes are transferred from the endosymbiont genome to the eukaryotic nucleus in a process called endosymbiotic gene transfer (EGT). These and other gene products, now under the control of host gene expression, are imported back into the endosymbiotic compartment to regulate endosymbiont growth and division. This definition has since been commonly applied^4,5^. However, the underlying hypothesis for organelle evolution — that genetic integration resulting from EGT and/or import of host-encoded gene products is essential for maintaining the endosymbiont as an integral cellular compartment — has not been rigorously tested.

In the decades since, increased sampling of eukaryotic diversity has uncovered evidence that, amongst microbes, endosymbioses are a common strategy for acquisition of new functions. Based on observations of EGT and host protein import, new organelles have been recognized: the chromatophore in *Paulinella chromatophora*^6–9^ and UCYN-A in *Braarudosphaera bigelowii*^10^. EGT and host protein import have also been observed in obligate, vertically-inherited nutritional endosymbionts of the parasite *Angomonas deanei* and insects, which are not formally recognized as organelles^11,12^. With the benefit of these newer models, our understanding of genetic integration has become more nuanced^13,14^. For example, the majority of host proteins imported into the *Paulinella* chromatophore do not originate from EGT but rather horizontal gene transfer (HGT) from other bacteria or eukaryotic genes^15^, showing that a host’s repertoire of pre-existing genes may play an outsized role in facilitating genetic integration^16^. UCYN-A was initially described as having an unstable relationship with its host as it is often lost from host cells during isolation and in culture, suggesting environmental conditions can affect endosymbiont stability despite genetic integration^17^. There have been bigger surprises: Some organisms temporarily acquire plastids from partially-digested prey algae^18,19^. The retained chloroplasts, called kleptoplasts, perform photosynthesis and, in several species, depend on imported host proteins to fill gaps in their metabolic pathways. Despite their genetic integration, these kleptoplasts cannot replicate in the host cell and are not required for host cell survival, indicating that genetic integration is not sufficient to achieve stable integration of the endosymbiont^20–22^. These findings highlight the importance of studying biodiverse organisms to inform new hypotheses for endosymbiotic evolution.

Amongst new model systems, *Epithemia* spp. diatoms offer a unique perspective on organellogenesis. These photosynthetic microalgae contain diazotroph endosymbionts (designated diazoplasts) that perform nitrogen fixation, a biological reaction that converts inert atmospheric nitrogen to bioavailable ammonia^23–28^. The ability to fix both carbon and nitrogen fulfills a unique niche in ecosystems. Numerous *Epithemia* species are globally widespread in freshwater habitats and have recently been isolated from marine environments^29–31^. The *Epithemia* endosymbiosis is very young relative to mitochondria and chloroplasts, having originated ∼35 Mya, based on fossil records^32^. Nonetheless, diazoplasts are obligate endosymbionts which are coordinately inherited during host cell division and present in all *Epithemia* species described so far, indicating co-evolution of diazoplasts and their host algae. Finally, *Epithemia* diazoplasts are closely related to UCYN-A, the diazotroph endosymbiont of *B. bigelowii* which was recently designated the first nitrogen-fixing organelle, or nitroplast^10,28,33^. Both *Epithemia* diazoplasts and UCYN-A evolved from free-living *Crocosphaera* cyanobacteria that have engaged in endosymbioses with several host microalgae. The independent evolution of free-living *Crocosphaera* into diazotroph endosymbionts in multiple host lineages enables comparisons that can lead to powerful insights.

If the significance of organelles lies in their function as integral cellular compartments, then metabolic and cellular integration with the host cell are paramount^34^. By these criteria, diazoplasts show a level of host-symbiont integration comparable to UCYN-A. Nitrogen fixation requires large amounts of ATP and reducing power, energy that can be supplied by photosynthesis. Yet nitrogenase, the enzyme that catalyzes nitrogen fixation, is exquisitely sensitive to oxygen produced during oxygenic photosynthesis. In free-living *Crocosphaera*, photosynthesis and nitrogen fixation is temporally separated such that fixed carbon from daytime photosynthesis is stored as glycogen to fuel exclusively nighttime nitrogen fixation. Diazoplasts have lost all photosystem genes and depend entirely on host photosynthesis for fixed carbon^27,35^. Recently, we showed that host and diazoplast metabolism are tightly coupled to support continuous nitrogenase activity throughout the day-night cycle in *E. clementina*: Diatom photosynthesis is required for daytime nitrogenase activity in the diazoplast, while nighttime nitrogenase activity also depends on diatom, rather than diazoplast, carbon stores^28^. In comparison, UCYN-A has lost only photosystem II and is dependent on both host photosynthesis and, likely, its own photosystem I, restricting it to daytime nitrogen fixation^36,37^. *Epithemia* spp. described so far typically contain 1-2 diazoplasts per cell that are vertically inherited during asexual cell division^25,28,30^. Diazoplasts have further been shown to be uniparentally inherited during sexual reproduction, similar to mitochondria and chloroplasts^38^. Coordinated replication of UCYN-A with host cell division has been observed to maintain a single endosymbiont per cell^10^. Similar mechanisms are likely in place to coordinate diazoplast inheritance with diatom division. In fact, the presence of diazoplasts in diverse *Epithemia* species globally widespread in freshwater and marine ecosystems demonstrates that the mechanisms of inheritance are robust through speciation events. Diazoplasts effectively serve as dedicated nitrogen-fixing compartments in *Epithemia*, whether or not genetic integration has occurred.

An important question emerges from these observations: is EGT and/or host protein import required to achieve the level of host-symbiont integration observed between diazoplasts and host *Epithemia*? Based on the similarity of diazoplasts to UCYN-A, the assumption is yes. However, there is evidence that metabolite exchange via endosymbiont-encoded transporters^39^ and division coordinated by host proteins outside the endosymbiotic compartment^11,40,41^ could form a stable compartment without genetic integration. We previously established freshwater *E. clementina* as a laboratory model for functional studies and herein performed *de novo* assembly and annotation of its genome. The genome sequence for *E. pelagica*, a recently-discovered marine species, was publicly released by the Wellcome Sanger Institute^30,42^. To facilitate comparison between these species, we also performed *de novo* genome annotation of *E. pelagica*. Notably, no genomes of *B. bigelowii* (which hosts UCYN-A) nor the eukaryotic host in any other diazotroph endosymbiosis have been available. We report genome and transcriptome analyses of these two *Epithemia* species as well as proteome analyses of *E. clementina* with the goals of 1) providing a necessary resource to accelerate investigation of this model and 2) elucidating the role of genetic integration in this very young, stably integrated endosymbiont.

## RESULTS

### The genome of *E. clementina* is larger and more repetitive than *E. pelagica*

*Epithemia* spp. are raphid, pennate diatoms composed of at least 50 freshwater species and 2 reported marine species^29,30^. Isolation and characterization of freshwater *E. clementina* was previously reported^28^ (Figure S1A). We isolated high molecular weight DNA from axenic *E. clementina* cultures,performed sequencing by long-read Nanopore and short-read Illumina and assembled a 418 Mbp haploid assembly with a high level of heterozygosity of 1.48% (Figure S1B). The final reported haploid assembly is complete, contiguous, and of high sequence quality (Figure S1C, Table 1). A chromosome-level 60 Mbp genome assembly (GCA_946965045) of *E. pelagica*, a marine species, was reported by the Sanger Institute^42^. Whole-genome alignments of *E. clementina* and *E. pelagica* did not show significant syntenic blocks in their nuclear genomes (Figure S1E). In contrast, their diazoplast genomes showed 5 major and 2 minor syntenic blocks (Figure S1F), similar to the synteny reported between diazoplast genomes of other *Epithemia* species^33,35^.

**Table 1.** *Epithemia* genome assembly statistics. Summary of assembly statistics for *E. clementina* and, where applicable, *E. pelagica*. Quality value (QV) represents a log-scaled estimate of the base accuracy across the genome, where a QV of 40 is 99.99% accurate. N50 and L90 are measures of genome contiguity. N50 represents the contig length (bp) such that 50% of the genome is contained in contigs ≥ N50. L90 represents the minimum number of contigs required to contain 90% of the genome. Finally, BUSCO (Benchmarking of Single Copy Orthologues) is an estimate of completeness of the genome (BUSCO_genome_) and proteome (BUSCO_protein_) of *E. clementina* and *E. pelagica*. ‘-‘ indicates no statistic.

|  | *E. clementina* | *E. pelagica* |
| --- | --- | --- |
| Genome size (bp) | 418,007,894 | 60,195,788 |
| GC | 44.3% | 48.19% |
| QV | 38.52 | - |
| Contig/chromosome # | 642 | 15 |
| N50 | 1,108,441 | - |
| L90 | 412 | - |
| Gene # | 26,453 | 20,203 |
| Repeat % | 80% | 27.36% |
| BUSCO<sub>genome</sub> | 100% | 100% |
| BUSCO<sub>protein</sub> | 99% | 94% |
| Diazoplast genome size (bp) | 3,072,807 | 2,483,960 |
| Diazoplast gene # | 1,910 | 1,679 |

Both *E. clementina* and *E. pelagica* genomes were annotated using evidence from protein orthology and transcriptome profiling. The nuclear genomes were predicted to contain 20,203 genes in *E. pelagica* and 26,453 genes in *E. clementina* (Figure 1A). The completeness of their predicted proteomes was assessed based on the presence of known single-copy orthologs in stramenopiles, yielding BUSCO_protein_ scores of 99% for *E. clementina* and 94% for *E. pelagica* (Table 1, Figure S1D). While the gene numbers between *E. clementina* and *E. pelagica* are similar and typical for diatoms, the genome of *E. clementina* is 7 times larger (Figures 1A and 1B, Table 1). The increased genome size is due to a substantial repeat expansion unique to *E. clementina* (Figure 1B and 1C). Notably, the differences in genome size observed amongst diatoms is largely due to repeat content (Figure 1B). Multiple LTR families and DNA transposons show expansions that contribute to the high repeat percent in *E. clementina* (Figure 1C, Table S2). This cross-family expansion may indicate a history of relaxed selection on the repeatome of *E. clementina*, perhaps related to a species bottleneck at the transition to freshwater habitats.

**Figure 1.**
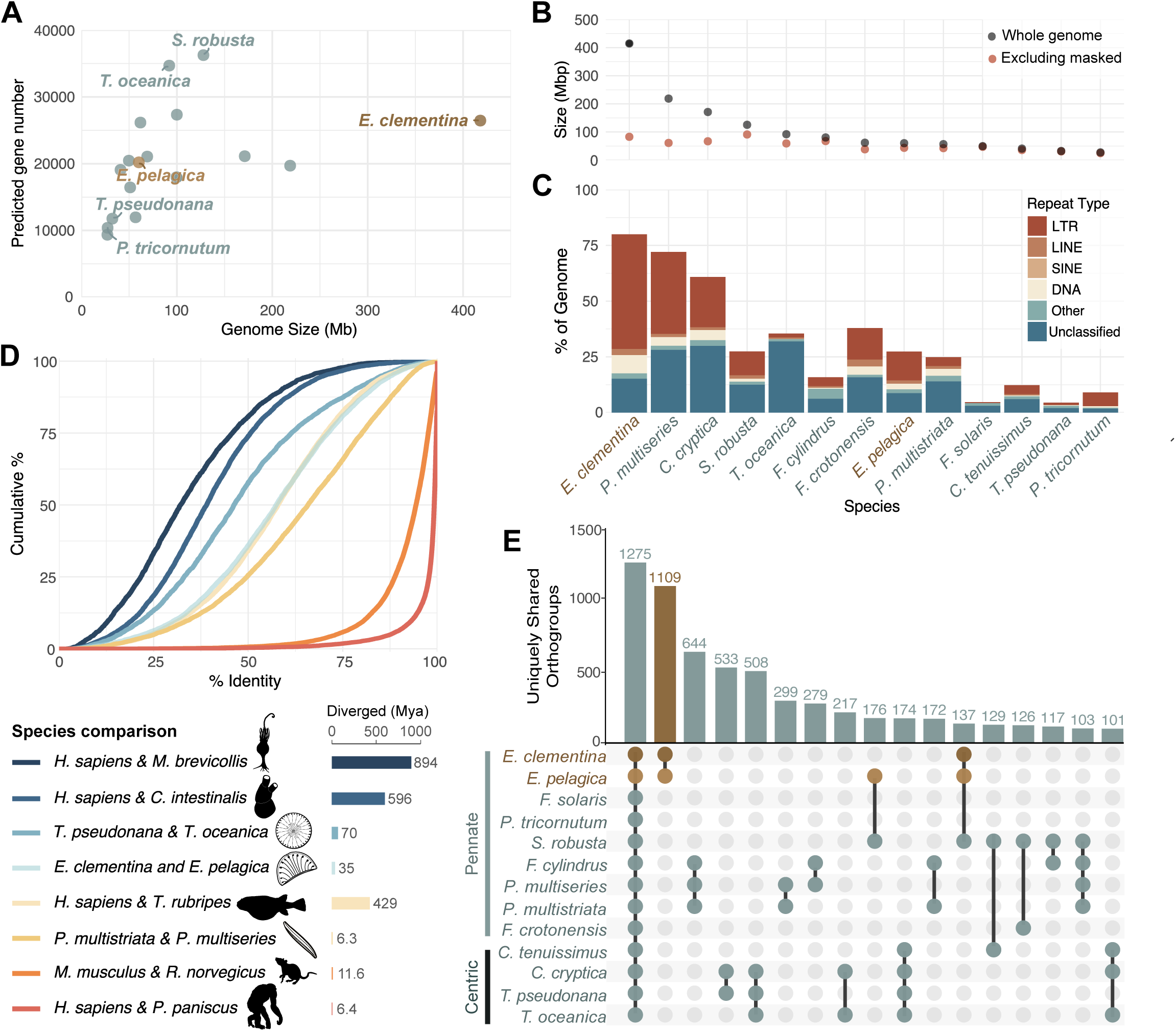
Highly divergent *E. clementina and E. pelagica* genomes share many unique gene families. **(A)** Genome size and total gene number for published diatom genomes compared with *Epithemia* species (dark blue). (See also Figure S1, Table S1.) **(B)** Comparison of repeat content in diatom genomes showing size of the whole genome (grey dots) or the genome excluding repeat elements (orange dots). X-axis is the same as 1C. **(C)** Breakdown of repeat types in diatom genomes showing amount in Mbp of the genome occupied by repeat elements of specific class, indicated by color. (See also Table S2.) **(D)** Cumulative distribution of amino acid identity between pairwise orthologs from reference species. Estimated divergence time of species pair is indicated (right bar graph). (See also Figure S2.) **(E)** UpSet plot depicting the number of uniquely shared orthogroups between all diatom species (first column) or subsets of 2-4 species. Orthogroups shared by *E. pelagica* and *E. clementina* are highlighted in brown. Columns are ranked by the number of uniquely shared orthogroups. (See also Figure S2.)

### Highly divergent *E. clementina and E. pelagica* genomes share many unique gene families

As a measure of divergence at a functional level, we compared the amino acid identity between orthologs across proteomes of several pairs of representative diatom and metazoan species (Figure 1D). Despite their estimated 35 Mya of speciation, *E. pelagica* and *E. clementina* showed a similar distribution of identity across protein orthologs as humans and pufferfish (*Homo sapiens* and *Takifugu rubripes*), which are estimated to have shared a common ancestor 429 Mya^43^. This rapid divergence, relative to age, is also observed in other diatom species, for example, *T. pseudonada*/ *T. oceanica* (70 Mya) and *P. multistriata*/ *P. multiseries* (6.3 Mya)^43^ (Figure 1D and Figure S2A). The loss of synteny, substantial differential repeat expansion, and low protein ortholog identity suggest that *E. pelagica* and *E. clementina* have diverged substantially during speciation, reflecting the rapid evolution rates of diatoms^44,45^ (Figure 1D and Figure S2A).

Because rapid divergence is common across diatoms, we evaluated the gene content of *E. pelagica* and *E. clementina* in comparison with other diatom species with complete genomes available. Gene families, defined by orthogroups, were identified for each species. We then quantified the number of uniquely shared gene families between subsets of diatom species, i.e. gene families shared between that group of species and not found in any other diatoms. Of 10,740 and 10,612 gene families identified in *E. clementina* and *E. pelagica* respectively, they share 8,942 gene families, a greater overlap than is observed between any other pair of diatom species (Figures S2A). Of these, 1,109 gene families are uniquely shared between *E. clementina* and *E. pelagica*, more than any other species grouping including the more recently speciated *Pseudo-nitzchia* species (Figure 1E). This *Epithemia-*specific gene set is significantly enriched for functions relating to carbohydrate transport and membrane biogenesis, which may have been important for adaptation to the endosymbiont (Figure S2B).

Because HGT is known to be a source of genes for endosymbiont functions expressed by the host^9^ and 3-5% of diatom proteomes have been attributed to bacterial HGT^46^, a significantly greater proportion than detected in other eukaryotic proteomes^47^, we identified candidate HGTs in the *Epithemia* genomes. A total of 118 and 97 candidate HGTs were identified in *E. clementina* and *E. pelagica* respectively, of which 51 are only detected in *Epithemia* (Figures S2C, S2D and S2E, Table S3). Notably, the two *Epithemia* species share a greater number of predicted horizontally-acquired genes than other diatom species and more than expected by gene family overlap alone. However, we were unable to identify enriched functions in this relatively small gene set that were informative. Overall, the uniquely shared features of the divergent nuclear genomes of *Epithemia* genus are valuable for identifying potential signatures of endosymbiotic evolution.

### Diazoplast-to-nucleus transfer of DNA is actively occurring in *E. clementina*

Having broadly compared the *Epithemia* genomes for shared features, we turned to specifically interrogate genetic integration between *Epithemia* and their diazoplasts. We specifically distinguished EGT, which we defined as the transfer of functional genes from the endosymbiont to the host nucleus, from endosymbiont-to-nucleus transfers of DNA, which is believed to be more frequent and often nonfunctional. Indeed, it has been shown that nuclear integrations of organellar DNA originating from mitochondria (designated NUMT) and plastids (NUPT) still occur^48,49^. Given the significantly younger age of the diazoplast, it is not clear whether nuclear integrations of diazoplast DNA (which we will refer to as NUDT) and/or functional transfers of genes (EGT) have occurred. To identify transfers of endosymbiont DNA to the host nucleus, we performed homology searches against the nuclear assemblies of *E. clementina* and *E. pelagica.* As queries, we used the diazoplast genomes of 4 *Epithemia* species (including *E. clementina* and *E. pelagica*) and the 5 most closely-related free-living cyanobacteria species for which whole genomes were available (Table S1). To prevent spurious identifications, alignments were excluded if they were <500 contiguous base pairs in length. In *E. clementina*, we identified seven segments, ranging from 1700-6400bp, with homology to the *E. clementina* diazoplast (Figures 2A and 2B, Data S1A-S1G). No homology to free-living cyanobacteria genomes was detected. The *E. coli* genome and a reversed sequence of the *E. pelagica* diazoplast were used as negative control queries and yielded no alignments. Finally, no regions of homology to any of the queries were detected in the nuclear genome of *E. pelagica*.

**Figure 2.**
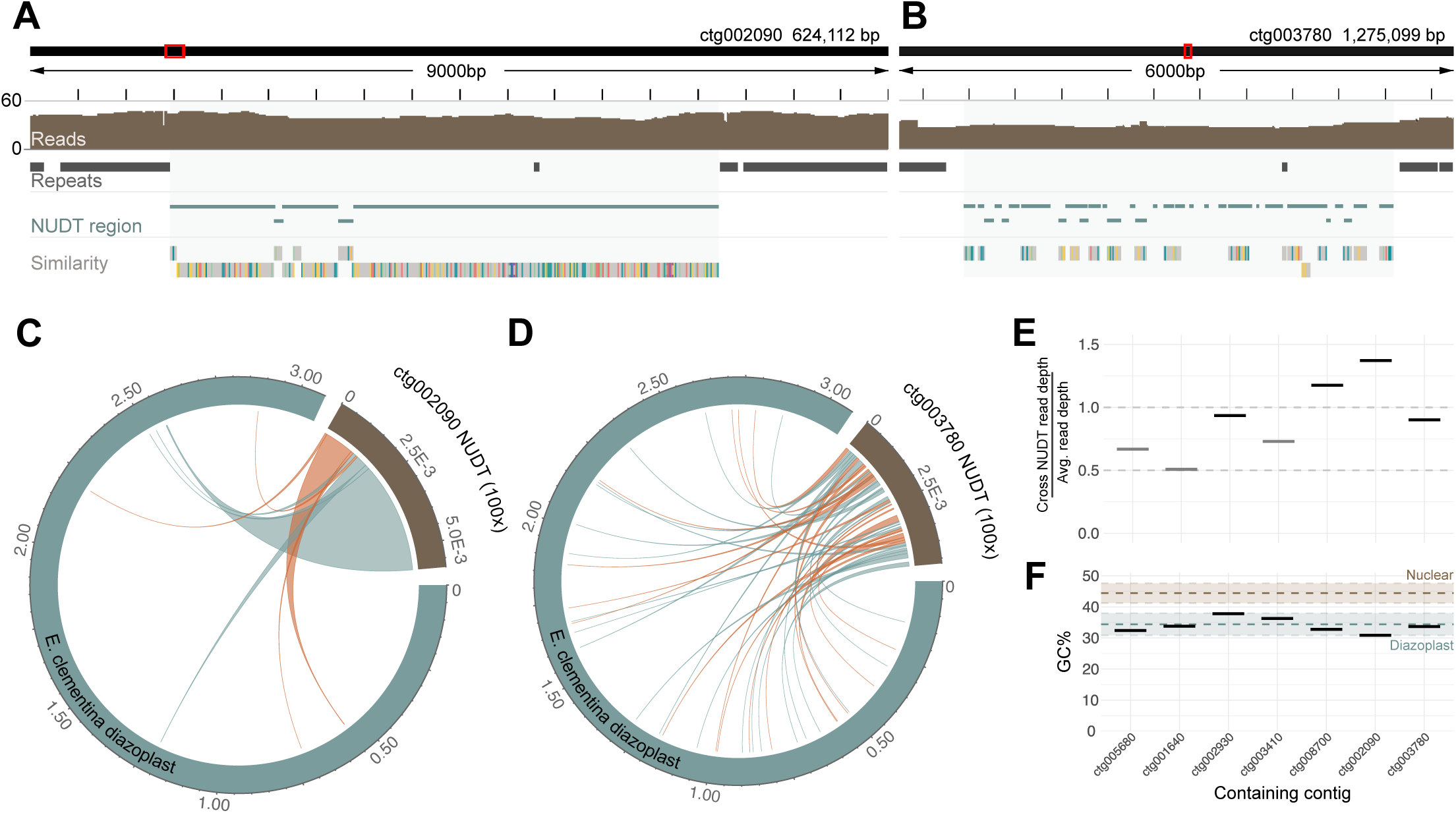
Detection of nuclear integrations of diazoplast DNA (NUDTs) **(A)** A representative, NUDT containing *E. clementina* nuclear genome locus on contig ctg002090. Tracks shown from top to bottom: nuclear sub-region being viewed (red box) within the contig (black rectangle); length of the sub-region, with ticks every 500bp; nanopore sequencing read pileup, showing long read support across the NUDT; location of repeat masked regions (dark grey bars); locations of homology to *E. clementina* diazoplast identified by BLAST, demarcating the NUDT (blue shade); regions of homology to the *E. clementina* diazoplast identified by minimap2 alignment, colors represent SNVs between the diazoplast and nuclear sequence. (See also, Data S1F.) **(B)** Same as A, for the NUDT on contig ctg003780. (See also, Data S1G.) **(C)** Circlize plot depicting the fragmentation and rearrangement of NUDTs. The diazoplast genome (blue) and the NUDT on contig ctg002090 (brown) with chords connecting source diazoplast regions to their corresponding nuclear region, inversions in red. The length of the NUDT is depicted at 100x true relative length for ease of visualization. (See also, Figure S3A-S3E.) **(D)** Same as C, for the NUDT on contig ctg003780. **(E)** Ratio of long read depth of NUDT compared to average read depth for the containing contig. Heterozygous insertions (light grey bars) show approximately 0.5x depth; homozygous insertions (black bars) show approximately 1.0x depth. **(F)** GC content of NUDTs, compared to mean GC content for 5kb sliding windows of the diazoplast genome (blue dashed line) and the nuclear genome (brown dashed line). Shaded regions represent mean ± 1 SD.

NUDTs showed features suggesting they were distinct from diazoplast genomic sequences and unlikely to be assembly errors. First, 4 of the 7 NUDTs were supported by long reads equivalent to 1x coverage of the genome indicating the insertions were homozygous. 3 NUDTs contained on ctg003410, ctg001640, ctg005680 showed the equivalent of 0.5x genome coverage, consistent with a heterozygous insertion in the diploid eukaryotic genome (Figure 2E, Data S1A, S1B, and S1D). Second, NUDTs had low GC content similar to that of the diazoplast but contain many single nucleotide variants (SNVs) with mean identity of 98.4% to their source sequences, indicative of either neutral or relaxed selection (Figures 2F and 3B). Finally, each NUDT was composed of multiple fragments corresponding to distal regions in the endosymbiont genome, ranging from as few as 8 distal fragments composing the NUDT on ctg002090 to as many as 42 on ctg003780 (Figures 2C, 2D, and S3A-S3E). This composition of NUDTs indicates either that fragmentation and rearrangement of the diazoplast genome occurred prior to insertion into the eukaryotic genome or that NUDTs were initially large insertions that then underwent deletion and recombination. Overall, the detection of NUDTs shows that diazoplast-to-nucleus DNA transfer is occurring in this very young endosymbiosis.

**Figure 3.**
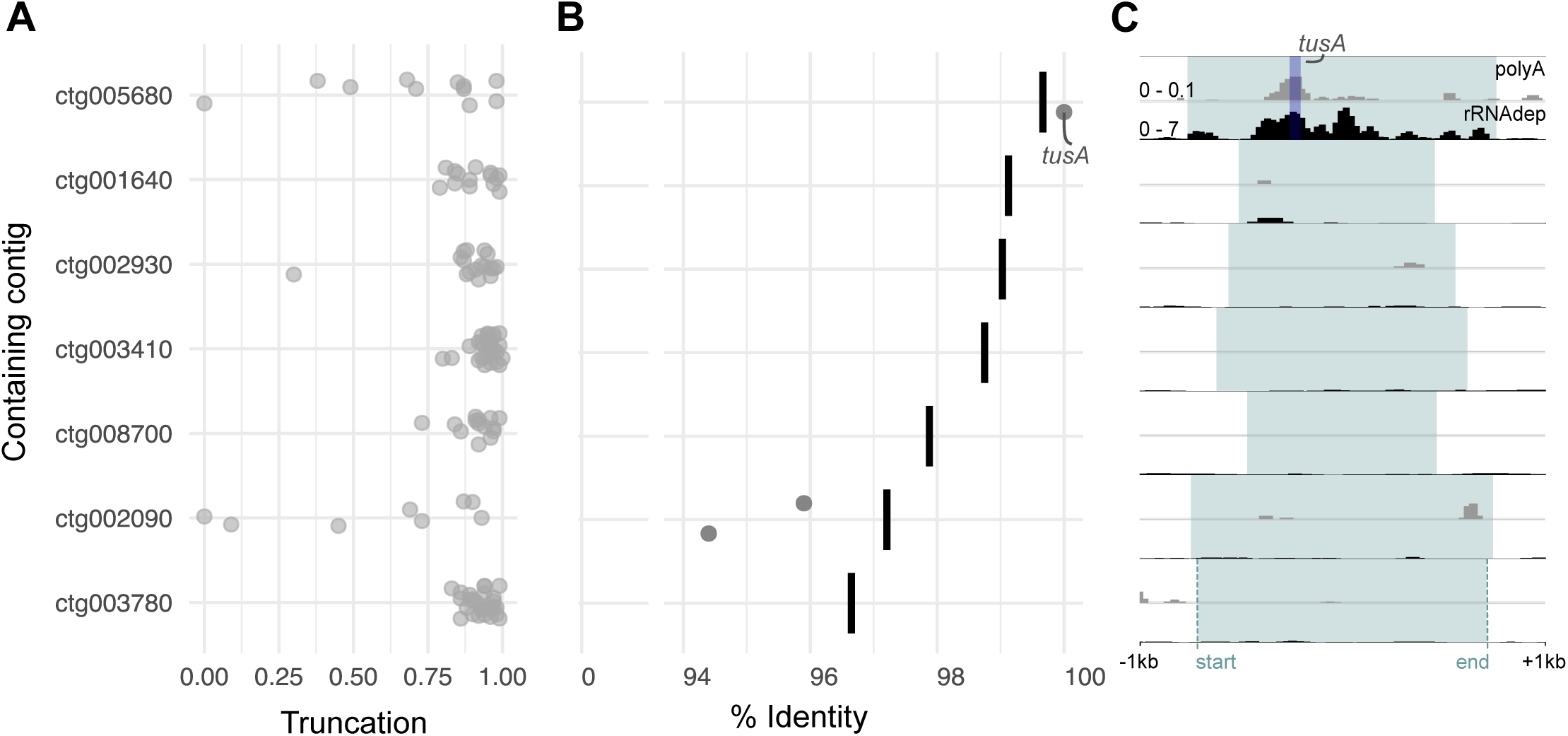
Most NUDTs are decaying and non-functional. **(A)** Truncation of diazoplast genes contained within each NUDT relative to the full-length diazoplast gene. **(B)** Nucleotide identity of diazoplast genes that are <30% truncated (points) contained within each NUDT compared to identity of the full containing NUDT sequence (bars). (See also, Figures S3F and S3G.) **(C)** Normalized expression across each NUDT (blue highlight) +/- 1kb of the genomic region surrounding the NUDT. For each NUDT, a pair of tracks shows RNA-seq reads after polyA enrichment of whole RNA plotted within background signal range, from 0 - 0.1 BPM (top, grey) and RNA-seq reads after rRNA depletion of whole RNA, plotted from 0 - 7 BPM (bottom, black). The region corresponding to the *tusA* gene in ctg005680 is highlighted in dark blue (See also Data S1A-S1G.)

### Most NUDTs are decaying and non-functional

To determine whether any of the identified NUDTs have resulted in EGT, we identified diazoplast genes present in NUDTs and evaluated their potential for function. A total of 124 diazoplast genes and gene fragments were carried over into the NUDTs (Figure 3A). (A few of these diazoplast genes have conserved eukaryotic homologs and were also predicted as eukaryotic genes in the *E. clementina* genome annotation (Data S1A-S1G).) 121 diazoplast genes detected in NUDTs are truncated >30% compared to the full-length diazoplast gene (Figure 3A). Of the three remaining, two genes contained on ctg002090 showed accumulation of SNVs that resulted in a premature stop codon and a nonstop mutation (Figures 3B, S3F and S3G). We performed transcriptomics to assess the expression from NUDTs. Neither of the two genes on ctg002090 showed appreciable expression. All except one NUDT showed <2 bins per million mapped reads (BPM), equivalent to background transcription levels within the region (Figures 3C and Data S1A-S1G). The truncation, mutation accumulation, and lack of appreciable expression of diazoplast genes encoded in NUDTs suggest that most are nonfunctional.

Only a single EGT candidate was detected contained on ctg005680: an intact sulfotransferase (*tusA*) gene that is 100% identical to the diazoplast-encoded gene (Figures 3A and 3B, Data S1A). The NUDT that contains this candidate appears to be very recent as it is heterozygous and shows 99.7% identity to the source diazoplast sequence (Figure 3B). Interestingly, *tusA* is implicated in Fe-S cluster regulation that could be relevant for nitrogenase function. Due to the high sequence identity, it is not possible to distinguish transferred *tusA* from that of *tusA* encoded in the diazoplast genome by sequence alone. However, transcript abundance above background levels was only detected in rRNA depleted samples that contain diazoplast transcripts and not in polyA-selected samples that remove diazoplast transcripts, indicating that the observed expression is largely attributed to diazoplast-encoded *tusA* (Figure 3C). Moreover, host proteins imported to endosymbiotic compartments often use *N*-terminal (occasionally *C*-terminal) targeting sequences^10,50,51^. We were unable to identify any added sequences in the transferred *tusA* indicative of a targeting sequence; the sequence immediately surrounding consisted only of native diazoplast sequence carried over with the larger fragment (Data S1A). Though there is no evidence for gene function, the transfer of this intact gene indicates that the conditions for EGT are present in *E. clementina*^4^.

### Few host-encoded proteins are detected in the diazoplast proteome

The critical step in achieving genetic integration is evolution of pathways for importing host proteins into the endosymbiont. While EGT and HGT from other bacteria can expand the host’s genetic repertoire, neither transferred genes nor native eukaryotic genes can substitute for or regulate endosymbiont functions unless the gene products are targeted to the endosymbiotic compartment. Abundant host-encoded proteins were detected in the proteomes of recently-acquired endosymbionts that have been designated organelles: 450 in the chromatophore of *Paulinella*^9^ and 368 in UCYN-A^10^. In both the chromatophore and UCYN-A, several host-encoded proteins detected in the endosymbiont fulfill missing functions that complete endosymbiont metabolic pathways, providing further support for the import of host-encoded proteins.

To determine whether host protein import is occurring in the diazoplast, we identified the proteome of the *E. clementina* diazoplast. We were unable to maintain long-term *E. pelagica* cultures to perform proteomics for comparison. Diazoplasts were isolated from *E. clementina* cells by density gradient centrifugation (Figure 4A). The purity of isolated diazoplasts was evaluated by light microscopy. The protein content of isolated diazoplasts and whole *E. clementina* cells containing diazoplasts were determined by LC-MS/MS. A total of 2481 proteins were identified with ≥2 unique peptides: 754 proteins were encoded by the diazoplast genome (detected/total protein coding = 43% coverage) and 1727 proteins encoded by the nuclear genome (6.5% coverage) (Figure 4B, Table S4). Of note, TusA, the only EGT candidate identified, was not detected in either proteome. To identify proteins enriched in either the diazoplast or host compartments, we compared protein abundance in isolated diazoplasts and whole cell samples across 3 biological replicates (Figure 4C). 492 diazoplast-encoded proteins were significantly enriched in the diazoplast and none were enriched in whole cell samples, supporting the purity of the isolated diazoplast sample. Similarly, most host-encoded proteins (1281) were significantly enriched in whole cell samples, indicating localization in host compartments. Six unique host-encoded proteins were significantly enriched in diazoplast samples, suggesting possible localization to the diazoplast. Five were encoded by Ec_g00815, Ec_g12982, Ec_g13000, Ec_g13118, and Ec_g25610. The sixth protein was encoded by two identical genes, Ec_g24166 and Ec_g03819, resulting from an apparent short duplication of it and two neighboring genes. Because the duplication makes Ec_g24166 and Ec_g03819 indistinguishable by amino acid sequence, we considered them one import candidate. Of the 6 host protein import candidates, Ec_g25610 and Ec_g13000 were detected only in the diazoplast sample, while the rest were identified in both diazoplast and whole cell samples. We are unable to rule out the possibility of nonspecific enrichment since neither genetics nor immunofluorescence are available in *E. clementina* to further validate their protein localization.

**Figure 4.**
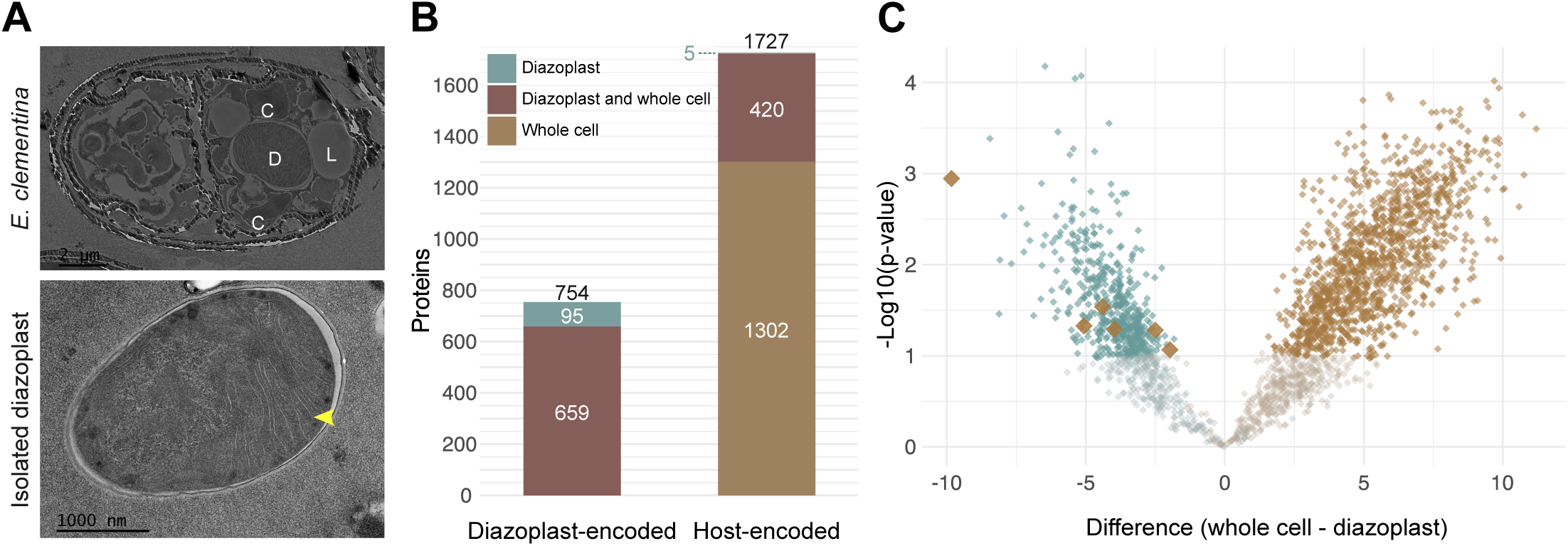
Few host-encoded proteins are detected in the diazoplast proteome. **(A)** Electron micrographs of (top) *E. clementina* cells with diazoplast (D), chloroplast lobes (C), and lipid bodies (L) indicated and (bottom) diazoplasts following purification with thylakoids (yellow arrow) indicated. **(B)** Number of diazoplast-encoded (left) and host-encoded (right) proteins identified by LC-MS/MS. Total number of proteins identified from each respective proteome is shown above each stacked bar. Colored bars and numbers indicate proteins identified in purified diazoplasts only, whole cells only, or both. **(C)** Volcano plot showing the enrichment of diazoplast-encoded (blue) and host-encoded (brown) proteins in whole cells or purified diazoplasts, represented by the difference between log2-transformed iBAQ values. Proteins enriched in the diazoplast are on the left side of the graph while those enriched in the host are on the right; the darker shade of each color represents significantly enriched hits. Host-encoded proteins significantly enriched in the diazoplast are shown with larger brown markers.

Instead we sought additional evidence to support the import of these host proteins by evaluating their potential functions^52^. No domains, GO terms, or BLAST hits (other than to hypothetical proteins found in other diatoms) were identified for any of the candidates except for Ec_g13118 which is annotated as an E3 ubiquitin ligase. In contrast to the unclear functions of these candidates, several host proteins detected in the chromatophore and UCYN-A proteome were assigned to conserved cyanobacterial growth, division, or metabolic pathways in these organelles. Moreover, none of the candidates for import into the diazoplast have apparent homology to proteins encoded in diazoplast or free-living *Crocosphaera* genomes to suggest they might fulfill unidentified cyanobacterial functions. Instead, all candidates are diatom-specific proteins: 3 candidates (Ec_g24166/Ec_g03819, Ec_g12982, and Ec_g13000) belonged to orthogroup OG0000250 which is uniquely shared with *E. pelagica* but no other diatoms (Figure 1E). The remaining 3 belonged to separate orthogroups (OG0001966, OG0004498, and OG0009247) which are shared broadly among diatoms including *E. pelagica*. Alignments of these candidates with their homologs did not show *N*- or *C*-terminal extensions consistent with possible targeting sequences for endosymbiont localization. Even if targeted to the diazoplast, we hypothesize that conserved eukaryotic proteins, especially diatom-specific proteins, are less likely to substitute for endosymbiont functions than bacterial proteins. Our functional analysis suggests these import candidates are unlikely to have critical functions in cyanobacterial pathways.

Finally, the detection of ∼100-fold fewer import candidates in the diazoplast indicate that host protein import, if occurring, is far less extensive than in the chromatophore or UCYN-A. Since the sensitivity of proteomics is highly dependent on biomass, we estimated the coverage of the diazoplast proteome based on the ratio of diazoplast-encoded proteins detected (754) compared to the total diazoplast protein-coding genes (1585). The coverage of the diazoplast proteome (48%) was comparable to the coverage of the published chromatophore proteome (422/867= 49%) and that of UCYN-A (609/1186= 51%) and therefore does not account for the low number of import candidates^9,10^. Overall, the number of host proteins detected in the diazoplast was significantly fewer and their functional significance unclear compared to host proteins detected in the chromatophore and UCYN-A.

## DISCUSSION

EGT and host protein import have been held as a necessary to achieve the “permanent” integration of organelles^2,5^. But this hypothesis for organelle evolution has been challenged by findings in young endosymbionts from diverse organisms^13,14,34^. We report analysis of two genomes of *Epithemia* diatoms and evaluate the extent of their genetic integration with their nitrogen-fixing endosymbionts (diazoplasts), thereby adding this very young endosymbiosis to existing model systems that can elucidate the integration of two cells into one.

### A window into the early dynamics of nuclear gene transfers

Our first significant finding was the detection of active diazoplast-to-nucleus DNA transfers but, as yet, no functional EGT in *Epithemia*. Our observations support findings in the chromatophore and UCYN-A that EGT is not necessary for genetic integration^10,15,53^. Given that EGT does not necessarily precede evolution of host protein import pathways, it may be a suboptimal solution for the inevitable genome decay in small asexual endosymbiont populations as a consequence of Muller’s ratchet^54,55^. Instead, the decayed nature of the NUDTs we detected in *E. clementina* is consistent with stochastic, transient, ongoing DNA transfer. Nonfunctional DNA transfers were previously only described from mitochondria or plastids with far more reduced genomes. The status of nuclear transfers from more recently-acquired organelles is unknown, as only protein-coding regions were used as queries to identify chromatophore transfers in *Paulinella* and only a transcriptome is available for the UCYN-A host, *B. bigelowii*^10,15,56^. NUDTs in *Epithemia* genomes therefore provide a rare window into the early dynamics of DNA transfer. For example, using the same homology criteria, we identified 5 NUMTs but no NUPTs in *E. clementina*. The NUMTs were significantly shorter than NUDTs and did not show rearrangement, which may suggest different mechanisms of transfer for NUDTs, NUMTs, and NUPTs in the same host nucleus. In addition, between-species differences may identify factors that affect transfer rates. The lack of observed NUDTs in *E. pelagica* suggest constraints on diazoplast-to-nucleus transfers. Previous observations in plant chloroplasts supported the limited transfer window hypothesis, which proposes the mechanism of gene transfer requires endosymbiont lysis and therefore the frequency of gene transfers correlates with the number of endosymbionts per cell^57–59^. However since *E. pelagica* and *E. clementina* contain similar numbers of diazoplasts per cell (1-2)^28,30^, the limited transfer window hypothesis does not explain the observed differences. Instead, there may be additional constraints imposed such as lower tolerance to DNA insertions in the comparatively smaller, non-repetitive genome of *E. pelagica*. Finally, the lack of NUDT gene expression, even with transfer of a full-length unmutated *tusA* gene, points to barriers to achieving eukaryotic expression from bacterial gene sequence. *Epithemia* is an apt model system to interrogate how horizontal gene transfer impacts eukaryotic genome evolution with at least 20 species easily obtained from freshwater globally and consistently adaptable to laboratory cultures^28–30,33,35^.

### Epithemia diazoplasts as a counterpoint to existing models of organellogenesis

A second unexpected finding was the detection of only 6 host protein candidates in the diazoplast proteome, much fewer and with less clear functional significance than in comparable endosymbionts that have been designated organelles. Methods for validating the localization of these import candidates are unavailable in *Epithemia.* Even if confirmed to target to the diazoplast, the candidates lack conserved domains or homology with cyanobacterial proteins to indicate they replace or supplement diazoplast metabolic function, growth, or division. Our findings are not explained by current models of organellogenesis that propose import of host proteins as a necessary step to establish an integral endosymbiotic compartment. In the traditional model described in the introduction, host protein targeting is a “late” bottleneck step required for the regulation of the endosymbiont growth and division. More recently, “targeting-early” has been proposed to account for establishment of protein import pathways prior to cellular integration as observed in kleptoplasts^19,20^. In this model, protein import is selected over successive transient endosymbioses, possibly driven by the host’s need to export metabolites from the endosymbiont via transporters or related mechanisms^60^. The establishment of protein import pathways then facilitates endosymbiont gene loss with metabolic functions fulfilled by host proteins leading to endosymbiont fixation. Contradicting both models, we observed minimal evidence for genetic integration despite millions of years of co-evolution resulting in diverse *Epithemia* species retaining diazoplasts, indicating that genetic integration is not necessary for its stable maintenance. At a minimum, the unclear functions of the few host proteins identified in the diazoplast proteome, if imported, suggest that the genesis of host protein import in *Epithemia* is very different than would be predicted by current models.

Diazotroph endosymbioses are fundamentally different from photosynthetic endosymbioses that are the basis for current organellogenesis models. First, the diazoplast is derived from a cyanobacterium that became heterotrophic by way of losing its photosynthetic apparatus. Regulation of endosymbiont growth and division by the availability of host sugars (without requiring an additional layer of regulation via import of host metabolic enzymes) may be more facile with heterotrophic endosymbionts maintained for a nonphotosynthetic function compared to autotrophic endosymbionts. It will be interesting to see how integration of the diazoplast differs from the endosymbiont of *Climacodium freunfeldianum*, another diazotrophic endosymbiont descended from *Crocosphaera* that likely retains photosynthesis^61,62^. Second, ammonia, the host-beneficial metabolite in diazotroph endosymbioses, can diffuse through membranes in its neutral form and does not require host transporters for efficient trafficking^63^. Previously, we observed efficient distribution of fixed nitrogen from diazoplasts into host compartments following ^15^N_2_ labeling^28^. Ammonia diffusion may have reduced early selection pressure for host protein import as posited by the targeting-early model. Finally, the eukaryotic hosts in most diazotroph endosymbioses are already photosynthetic, in contrast to largely heterotrophic hosts that acquired photosynthesis by endosymbiosis. For instance, cellular processes that enabled intracellular bacteria to take up residence in the ancestor of *Epithemia* spp. were likely different than those of the bacterivore amoeba ancestor of *Paulinella chromatophora*. Autotrophy and lack of digestive pathways would reduce the frequency by which bacteria might gain access to the host cell, such that the selection of host protein import pathways over successive transient interactions would be less effective. Overall, a universal model of organellogenesis is premature given the limited types of interaction that have been investigated in depth, highlighting instead the importance of increasing the diversity of systems studied.

### Are diazoplasts “organelles”?

As detailed in the introduction, diazoplasts show metabolic and cellular integration with their host alga comparable to that of UCYN-A, the first documented nitrogen-fixing organelle^10,17^. However, while hundreds of host proteins were detected in the UCYN-A proteome, including many likely to fill gaps in its metabolic pathways, a handful of host proteins with unknown function were detected in the diazoplast proteome. Based on the conventional definition which specifies genetic integration as the dividing line between endosymbionts and organelles, diazoplasts would not qualify^2,5^. However, over a decade ago, Keeling and Archibald^34^ suggested that “if we use genetic integration as the defining feature of an organelle, we will never be able to compare different routes to organellogenesis because we have artificially predefined a single route.” They further hypothesized that if an endosymbiont became fixed in its host absent genetic integration, “it might prove to be even more interesting… by focusing on how it did integrate, perhaps we will find a truly parallel pathway for the integration of two cells.” The diazoplast appears to be such a parallel case in which non-genetic interactions were sufficient to integrate two cells.

If not gene transfer and host protein import, then what is the “glue” that holds this endosymbiosis together? The loss of cyanobacterial photosystems and dependence on host photosynthesis indicate that diazoplasts acquired new transporters for host sugars^28^. There are several examples of cyanobacteria that express genes for sugar transport^39,64,65^. Therefore, acquisition via horizontal gene transfer from another bacteria to the diazoplast ancestor, prior to the endosymbiosis, rather than targeting of a eukaryotic transporter post-endosymbiosis, seems more likely. Consistent with this hypothesis, potential transporters were not detected amongst host protein import candidates detected in UCYN-A^10^. In fact, mixotrophy may have facilitated the adoption of an endosymbiotic lifestyle by the free-living cyanobacterium. Regardless of the timing of acquisition, sugar transport function could allow diazoplast growth to be regulated by nutrient availability from the host. Similarly, cytosolic host proteins may coordinate diazoplast division from outside the endosymbiotic compartment^11,41^. Eukaryotic dynamins required for mitochondria and chloroplast fission localize to the surface of the organellar outer membrane, acting coordinately with bacterial fission factors located in the organelle^40,66^. Diazoplasts appear to be surrounded by a host-derived membrane^26,67^ (which may be lost during diazoplast purification) (Figure 4A). In analogy to dynamins, host proteins localized to this outer membrane may mediate diazoplast fission without requiring protein import pathways. Finally, cell density^68^ and mechanical confinement^69^ have been demonstrated to limit the growth of cyanobacteria, suggesting that host regulation of the volume of the endosymbiotic compartment could also be an effective mechanism. The mechanisms for the robust metabolic exchange and coordinated division observed in diazoplasts will be the focus of future studies.

### Application of cell evolution models to bioengineering

Diazoplasts provide another example that the current organelle definition does not account for observations in many biological systems and may be overdue for revision to reflect biological significance in the spectrum of endosymbiotic interactions. At a minimum, it is time to disentangle the current definition of an organelle from models that elucidate the formation of integral cellular compartments. Identifying mechanisms to integrate cells is more than an academic exercise. The ability to engineer bacteria as membrane compartments to introduce new metabolic functions in eukaryotes would be transformative^70,71^. For example, nitrogen-fixing crop plants that could replace nitrogen fertilizers is a major goal for sustainable agriculture. Efforts to transfer the genes for nitrogen fixation to plant cells have been slow, hampered by the many genes required as well as the complex assembly, high energy requirements, and oxygen sensitivity of the reaction. We previously proposed an alternative strategy inspired by diazotroph endosymbioses: introducing nitrogen-fixing bacteria into plant cells as an integral organelle-like compartment^28^. This approach has the advantage that diazotrophs express all required genes with intact regulation, coupled to respiration, and in a protected compartment. Diazoplasts, which achieve stable integration without significant genetic integration, are an important alternative to UCYN-A and other organelles, which are defined by their genetic integration, to inform this strategy. Identifying the nongenetic interactions that facilitated diazoplast integration with *Epithemia* will be critical for guiding bioengineering.

### Ongoing genome reduction may drive genetic integration in diazotroph endosymbioses

The fewer number of host protein candidates and their lack of clear function in diazoplasts versus UCYN-A is not associated with differences in their function as nitrogen-fixing cellular compartments. Rather, an alternative explanation points towards differences in the extent of genome reduction in diazoplasts, which encode 1585-1848 protein-coding genes, compared to UCYN-A, which encode 1200-1246 protein-coding genes^72^. Among the genes missing from the UCYN-A genome are cyanobacterial IspD, ThrC, PGLS, and PyrE; for each, an imported host protein was identified that could substitute for the missing function^10^. In contrast, these genes are retained in diazoplast genomes, including those of *E. clementina* and *E. pelagica*, obviating the need for host proteins to fulfill their functions (Figure S4). Consistent with diazoplasts and UCYN-A being at different stages of genome reduction, diazoplast genomes contain >150 pseudogenes compared to 57 detected in the UCYN-A genome, suggesting diazoplasts are in a more active stage of genome reduction^27,33,35^. Interestingly, even genes retained in UCYN-A, namely PyrC and HemE, have imported host-encoded counterparts^10^. The endosymbiont copies may have acquired mutations resulting in reduced function, necessitating import of host proteins to compensate. Alternatively, once efficient host protein import pathways were established, import of redundant host proteins may render endosymbiont genes obsolete, further accelerating genome reduction. Genetic integration may in fact be destabilizing for an otherwise stably integrated endosymbiont, at least initially, as it substitutes essential endosymbiont genes with host-encoded proteins that may not be functionally equivalent and require energy-dependent import pathways. Comparing these related but independent diazotroph endosymbioses yields valuable insight, which otherwise would not be apparent. Diazoplasts at 35 Mya may represent an earlier stage of the same evolutionary path as ∼140 Mya UCYN-A, in which continued genome reduction will eventually select for protein import pathways. Alternatively, diazoplasts may have evolved unique solutions to combat destabilizing genome decay, for example through the early loss of mobile elements.^27,33,73^ Whether they represent an early intermediate destined for genetic integration or an alternative path, diazoplasts provide a valuable new perspective on endosymbiotic evolution.

### Limitations of the study

- Accurate gene family analysis is dependent on species sampling. While we sought to sample species representative of diatom diversity, some of our reported *Epithemia*-specific gene families may be shared by non-*Epithemia* species not present in the data set.
- While we included free-living *Crocosphaera* relatives for our homology search for NUDTs, we cannot eliminate the possibility that there are NUDTs or EGTs derived from sequences that were once in diazoplast genomes but have been lost.
- We did not observe expression associated with genes contained in NUDTs, however, we cannot exclude that they may be expressed under untested conditions.
- While differential centrifugation was an effective means of diazoplast enrichment, there may still be contamination by other cellular compartments.
- Low-abundance proteins may fall below the detection threshold in label-free proteomics. While our proteome coverage was comparable to previously reported studies, we cannot eliminate the possibility that there are undetected host proteins enriched in the diazoplast fraction.
- Tools such as immunofluorescence and genetics have not yet been established in *Epithemia* diatoms. We are therefore unable to confirm the localization or function of any *Epithemia* proteins.

## Supporting information

Table S1. SpeciesAccessions

Table S2. RepeatMaskerOutputs

Table S3. HGTGenes_EpithemiaSpecificOGs

Table S4. ProcessedProteomicsData

## RESOURCE AVAILABILITY

### Lead contact

Requests for further information and resources should be directed to and will be fulfilled by the lead contact, Ellen Yeh

### Materials availability

Cultures and reagents used in this study are available upon request from lead contact.

### Data and code availability

Data supporting the findings of this work will be made available upon publication on Mendeley at DOI:10.17632/rr9t3ccbc5.1. Sequencing data generated for this study will be made available at NCBI BioProject accession PRJNA1147773. The genome and annotation of *E. clementina* have been deposited and will be made available at NCBI accession [GCA ID TBD], TaxID 3042617.

The genome assembly pipeline is published at https://github.com/doenjon/Epithemia_assembly

Any additional information required to re-analyze the data reported in this paper is available from the lead contact upon request.

## ACKNOWLEDGEMENTS

We thank Chriz Schvarcz and Kelsey McBeain for the generous sharing of *E. pelagica* cultures. We thank Dr. Devaki Bhaya and Dr. Arthur Grossman and members of their labs for support and feedback on the project. We are grateful to Scott Miller and Heidi Abresch for their *Epithemia* expertise and discussion. Our thanks to Andres Reyes for mass spectrometry technical training. We thank Daniel S. Rokhsar, Jonathan Zehr, Kendra Turk-Kubo, Andy Alverson, Elizabeth Ruck, Paolo Carnevali, Dmitri Petrov, and Cedric Feschotte for advice during the project. Anti-NifDK polyclonal antibodies were kindly provided by Dennis Dean. E.Y. is a Chan-Zuckerberg Biohub – San Francisco Investigator and supported by Burroughs Wellcome Fund. S.F. was partially funded by NIH training grant (T32GM007276). M.S.O. was partially funded by NIH training grant (5T32AI007328-32). The contents of this manuscript are solely the responsibility of the authors and do not represent the official views of the NCRR or the National Institutes of Health. S.-L.X is funded by the NIH grant S10OD030441 and the Carnegie Endowment Fund to the Carnegie Mass Spectrometry Facility.

## AUTHOR CONTRIBUTIONS

S.F., M.S.O., and E.Y, wrote the manuscript with input from all authors. S.F. and E.Y. conceptualized project. S.F., J.D., M.S.O., T.B., and E.Y. contributed to project design and strategy. Initial DNA extraction and preliminary analysis was performed by S.F., J.D., and T.B. Axenic cultures were isolated by S.L.Y.M. Final DNA isolation and sequencing experiments were performed by S.F. and S.L.Y.M. S.F. isolated and performed RNA sequencing experiments. Final genome assembly was performed by J.D. Genome annotation and analysis were performed by S.F. S.-L.X. provided proteomics resources and expertise. M.S.O. performed and analyzed proteomics. All authors read and approved the final paper.

## DECLARATION OF GENERATIVE AI AND AI-ASSISTED TECHNOLOGIES

During the preparation of this work, the author(s) used GPT-4o mini and Claude 3.5 Sonnet for assistance with minor bioinformatic troubleshooting. The authors reviewed and edited all AI outputs and take full responsibility for the content of this publication.

## SUPPLEMENTAL INFORMATION

Supplemental Figures S1-S4

**Table S1.** NCBI Accessions and access links for all species used in this study, related to Figures 1, 2, S1 and S2.

**Table S2.** RepeatMasker output tables for *Epithemia*, related to Figures 1B and 1C

**Table S3.** Gene codes, putative identity, and gene tree of *Epithemia*-specific horizontally transferred genes, related to Figures 1D, 1E, and S2C-E.

**Table S4.** Processed and unprocessed proteomics data, related to Figure 4.

**Data S1.** Integrative genomics viewer screenshots of genomic context, expression, and read support of all NUDTs, related to Figures 1 and 2.

## SUPPLEMENTAL FIGURE TITLES AND LEGENDS

**Figure S1.**
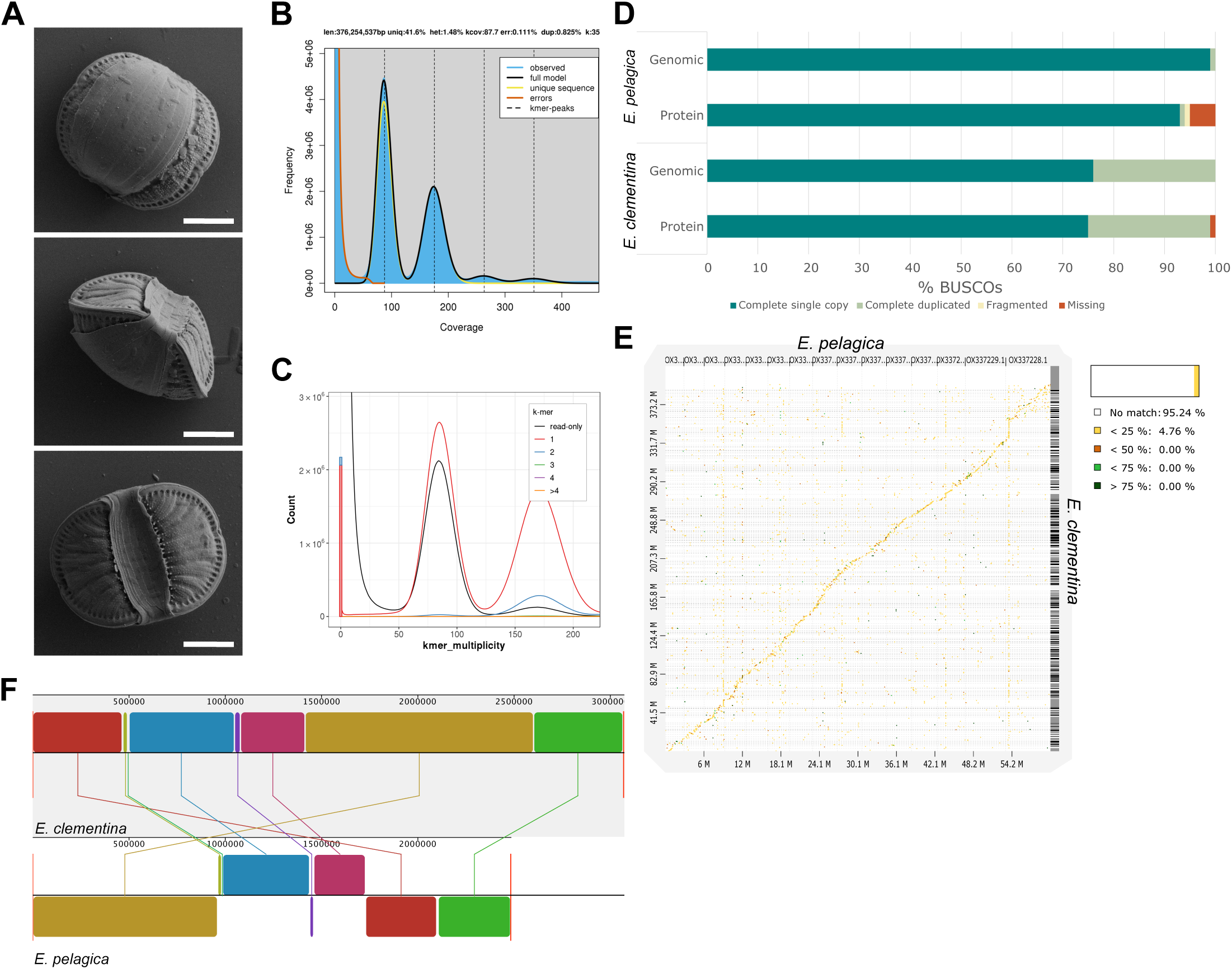
*E. clementina* genome assembly statistics and features. **(A)** Scanning electron micrographs of *E. clementina,* scale bar 5μm. Top: View looking down on the dorsal girdle band. Middle: View down the apical axis. Bottom: View of the ventral face, lined by prominent fenestral bars regularly spaced between the radial striae. The raphe lies along the strongly curved keel on the ventral margin and pinches slightly towards the dorsal margin. **(B)** GenomeScope spectrum of 35-mer multiplicity collected from the Illumina sequencing reads. Peak at 1x coverage (∼90) and 2x coverage (∼180), consistent with a diploid genome. **(C)** Merqury spectrum of k-mer multiplicity collected from the Illumina sequencing reads, stacked lines colored by number of times k-mer is seen in the genome assembly. Few k-mers within the heterozygous and homozygous peaks are read-only (black), suggesting that the assembly is not missing significant sequence present in the reads. **(D)** Stramenopile-specific Benchmarking Universal Single-Copy Orthologs (BUSCOs) for *E. pelagica* and *E. clementina* genomes and proteomes. Both genomes contain all stramenopile BUSCOs, however the *E. pelagica* annotation is less complete. The genome and proteome of *E. clementina* show some duplication. This duplication of stramenopile BUSCOs is similar or less than that found in previously published genomes of diatoms, for example, *Nitzschia putrida* (GCA_016586335, 39% duplicate BUSCOs)*, Fragilariopsis cylindrus* (GCA_900095095.1, 40% duplicate BUSCOs*), and Fragilaria radians* (GCA_900642245.1, 19% duplicate BUSCOs). **(E)** Whole genome alignment of the *E. clementina* and *E. pelagica* genome assemblies. White indicates no sequence homology, yellow indicates alignments at <25% nucleotide identity. There is only 4.76% sequence homology between the two genomes at the nucleotide level, all at <25% identity. **(F)** Genomic synteny between the whole genome alignments of the *E. clementina* and *E. pelagica* diazoplasts, showing 7 syntenic blocks.

**Figure S2.**
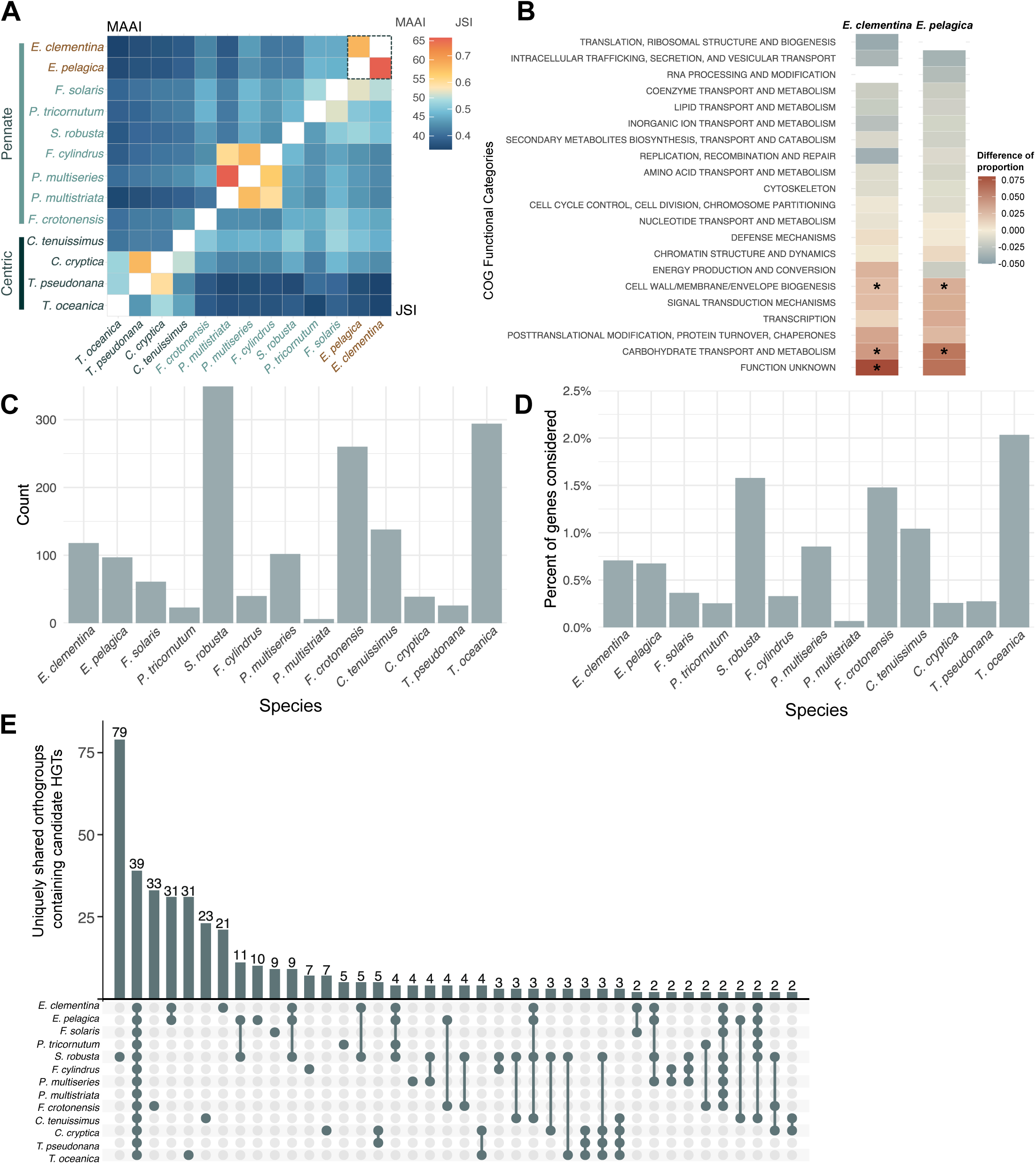
Gene family enrichment and HGT statistics. **(A)** Asymmetrical heatmap of ortholog comparisons between diatom species pairs, showing mean amino acid identity (MAAI) of pairwise orthologs (top) and Jaccard similarity index (JSI) of orthogroups (bottom). **(B)** Enrichment analysis of clusters of orthologous group (COG) categories in *E. clementina* and *E. pelagica.* Color indicates difference in proportion of each COG category within genes in *Epithemia* unique gene families and the whole functionally annotated gene set. An asterisk indicates categories that were found to be statistically significantly enriched. **(C)** Total count of HGT candidate genes across diatom species, considering only high-confidence candidates. **(D)** Proportion of the genes considered for HGT analysis that were identified as high-confidence HGT candidates. Genes were considered for analysis if they had >10 identifiable homologues. **(E)** UpSet plot depicting the number of uniquely shared, HGT candidate containing orthogroups between diatom species groupings (including singlets). Columns are ranked by the number of uniquely shared orthogroups.

**Figure S3.**
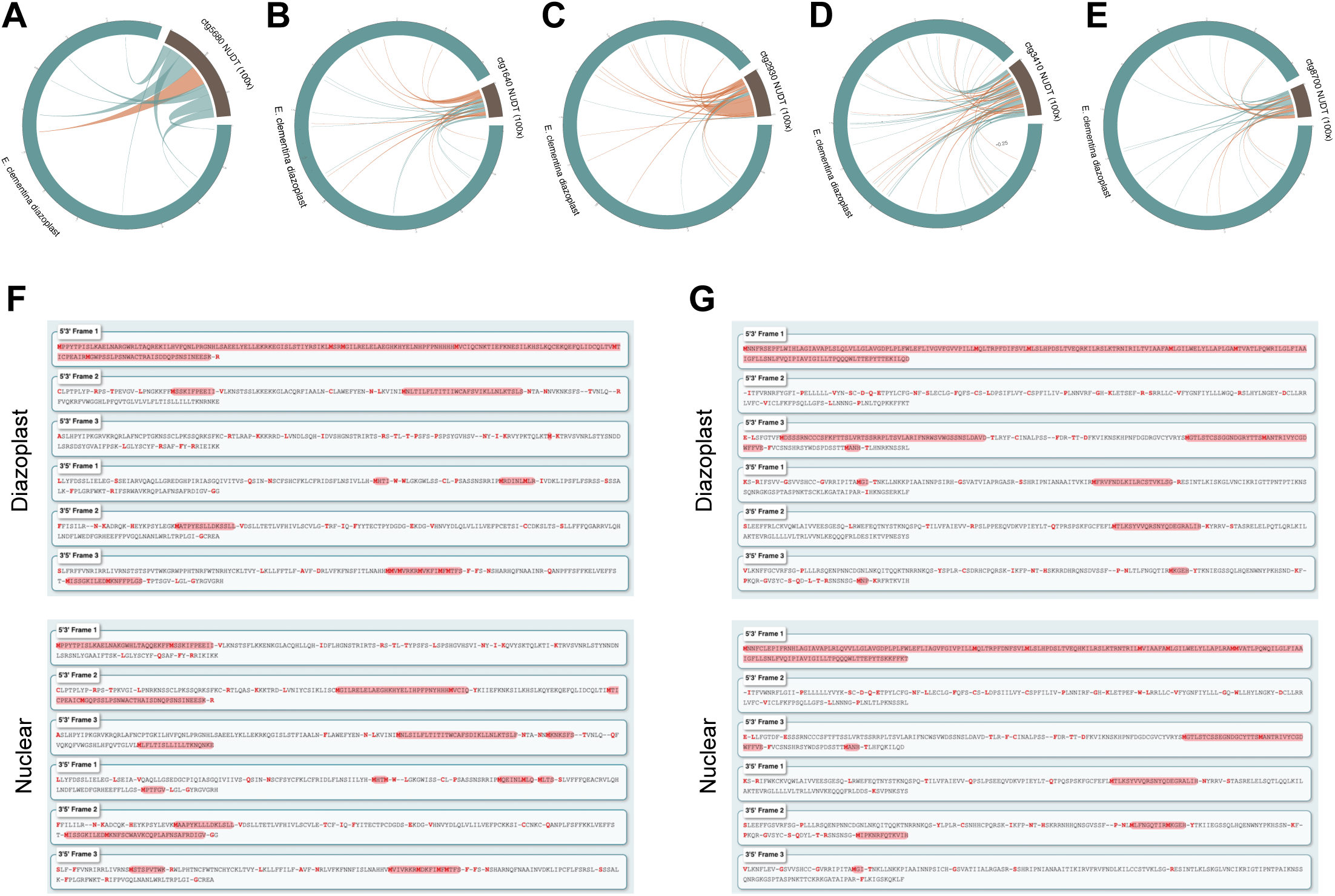
NUDT fragmentation and gene containing regions. **(A-E**) Circlize plots depicting the fragmentation and rearrangement of the NUDTs. The diazoplast genome (blue) and the NUDT on labeled contig (brown) with chords connecting source diazoplast regions to their corresponding nuclear region, inversions in red. The length of the NUDT is depicted at 100x true relative length for ease of visualization. **(F)** Translation in all potential frames of the gene contained within the NUDT on contig ctg002090 (transcriptional repressor, gene ID: P3f56_RS08570). The copy within the NUDT (bottom) is untruncated (100% of the full-length gene) by nucleotide sequence and is 96% identical to the corresponding diazoplast gene (top). Compared to the diazoplast-encoded gene, the gene contained in the NUDT has a mutation that results in a premature stop codon at amino acid 39 (out of 177). Red highlight indicates a potential translation. 5’3’ Frame 1 is the native diazoplast frame. **(G)** Translation in all potential frames of the gene contained within the NUDT on contig ctg002090 (low-complexity tail membrane protein, gene ID: P3F56_RS01750). The gene is 9% truncated at the 3’ terminus (91% of the full-length gene). Compared to the diazoplast-encoded gene, the gene contained in the NUDT has several non-synonymous mutations and is missing 16 amino acids at the C-terminus. Red highlight indicates a potential translation. 5’3’ Frame 1 is the native diazoplast frame.

**Figure S4.**
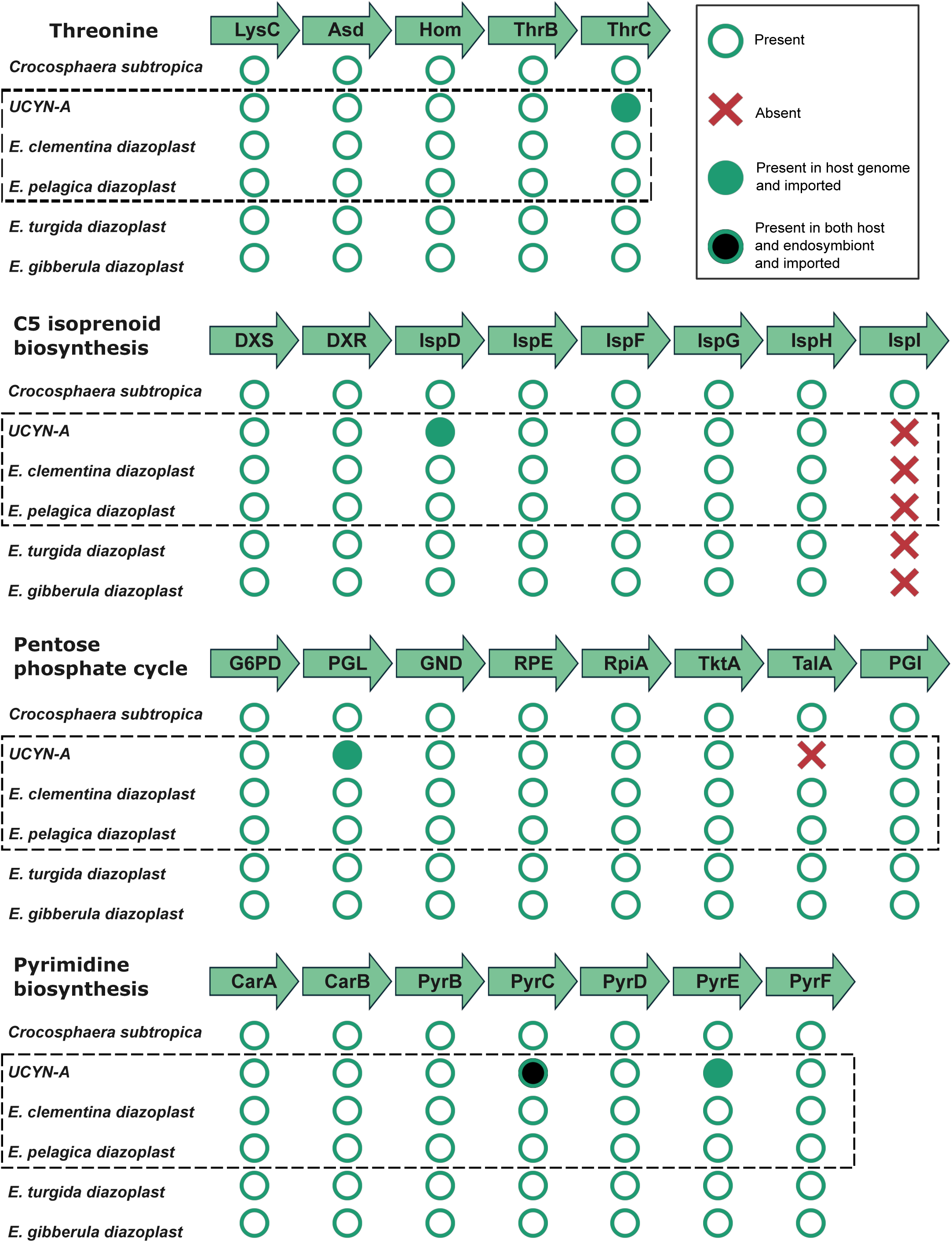
Comparative pathway analysis of diazoplasts and close relatives. KEGG pathway analysis of *E. clementina, E. turgida, E. gibberula* diazoplasts as well as *C. subtropica* and UCYN-A, indicating presence (green circle) or absence (red x) in the genome. Filled green circle indicates evidence for import of a host-encoded protein; filled black circle indicates presence in the endosymbiont genome and evidence for import of a host-encoded protein. A black dashed box indicates species for which data exists for both endosymbiont and host nuclear genes.

## SUPPLEMENTAL DATA TITLES AND LEGENDS

**Data S1: Detailed genome tracks across NUDT regions, related to Figures 3 and 4**

(**A-G**) For all NUDTs, full context genome tracks from the Integrated Genomics Viewer zoomed in to the NUDT region (left) or zoomed out to a 20kb surrounding region (right). Tracks from top to bottom are:

1) Region file of masked repeat regions;

2) Feature file of *E. clementina* gene models;

3) Region file of *E. clementina* diazoplast homology found by BLAST, demarcates the NUDT;

4-7) Alignment files of homology found by minimap2 when aligning 4) *E. clementina* diazoplast, 5) *E. pelagica* diazoplast, 6) *E. turgida* diazoplast, and 7) *E. gibberula* diazoplast to the *E. clementina* nuclear genome;

8-10) Normalized expression data in BPM of RNA seq from combined replicates of poly-adenylated transcript enriched RNA collected from three treatment conditions.;

11-13) Normalized expression data in BPM of RNA seq from combined replicates across of ribosomal RNA depleted RNA collected from three treatment conditions.;

14) Read pileup of axenic nanopore reads. Colored bars at certain sites indicate proportion of SNVs across the reads deviating from the haplotype reference assembly often resulting from a heterozygous site but sometimes from reads accumulating at undiscernible copies of repeat elements.;

15) Alignment file of axenic, nanopore long reads aligned to the reference *E. clementina* genome. An aligned read identical to the reference sequence is rendered as a single plain grey bar. Colors at sites along the read denote SNVs from the reference assembly. Small indels are denoted by small purple bars. A thin black bar within a read represents a region not present in the read that is present in the haplo-assembly (i.e. larger indels). Very light grey bars are secondary alignments, which accumulate at repeat elements.

## STAR★METHODS

### EXPERIMENTAL MODEL AND STUDY PARTICIPANT DETAILS

#### Diatom strains and culture conditions

Wild isolates of *E. clementina* were cultivated in CSi-N media in vented flasks under 10 μmole photon m - 2 s-1 of white light at 20°C. Cultivation procedures are also detailed in^28^. *E. pelagica* cells from wild isolates reported in Schvarcz et al.^30^ were shipped overnight from University of Hawaii at Manoa, HI in an insulated box. Upon arrival cultures were centrifuged at 1000 x g, media was aspirated, and cells were resuspended in trizol for RNA extraction.

### METHOD DETAILS

#### Generation of axenic culture

Cultures were initially isolated from a single cell of *E. clementina* and were thus clonal but xenic. To generate axenic cultures for sequencing, cells were incubated overnight with lysozyme to disrupt the cell walls of gram-positive bacteria, then treated to a 30-minute pulse of antibiotic cocktail (100 µg/mL Carbenicillin, 25 µg/mL Chloramphenicol, 5 mg/mL Levofloxacin, 50 mg/mL Rifampicin, 50 µg/mL Streptomycin). Immediately following, cultures were spray plated^74^. A small volume of dilute culture suspension was aspirated into a glass Pasteur pipette and held perpendicular to a stream of sterile air. The air atomizes the culture; the small droplets are then captured on a CSi-N agar plate. This process isolates single cells of *E. clementina* and disrupts their associated bacterial community. Cells on the agar plates were allowed to form colonies which were screened for any visual bacterial growth. Only those colonies that lacked bacterial growth were chosen for further cultivation. The resulting strain was expanded and confirmed axenic in subsequent sequencing experiments.

#### Scanning electron microscopy

Xenic cultures of *E. clementina* were resuspended, pelleted at 23°C at 1000 x g, rinsed with CSi-N media, and then resuspended in 250uL of PBS. Cells were transferred in a droplet to poly-L-lysine coated 12mm diameter glass cover slips and left to sit on a flat surface for 5min. The PBS was gently aspirated, and a droplet of 4% paraformaldehyde in PBS was added to coat the entire cover slip surface. Cells were fixed for 10min in the dark and then the cover slip was rinsed twice with PBS. An ethanol dehydration series was performed wherein cover slips were sequentially immersed in 60%, 70%, 80%, 90%, and 100% v/v ethanol in PBS. The cover slips were gently dried on a 42°C heat block. The cover slips were secured to a low-profile pin mount and sputter coated in a Leica ACE600 High Vacuum Sputter Coater with gold to a thickness of 6nm. Samples were imaged on a Zeiss Sigma FE-SEM.

#### High molecular weight DNA extraction

*E. clementina* were grown to a density of approximately 400,000 cells/mL and 20-30 million cells were used as input to HMW DNA extraction. Xenic cultures were first subjected to a round of centrifugation through a discontinuous Percoll gradient to deplete excess bacteria. *E. clementina* cells pellet out of the solution entirely, whereas a portion of their bacterial community stays suspended in various Percoll fractions. Centrifugation steps were performed at 23°C at 1000 x g. For both xenic and axenic cultures, HMW DNA was isolated using a nuclei extraction method^75^. Cells were suspended in a minimal volume of nuclear isolation buffer (NIB) and the transferred to a mortar where they were flash frozen and then ground with the pestle until a paste formed. This grinding process was repeated a total of three times. Cell homogenate was transferred to a 15mL falcon tube containing NIB, rinsing the mortar with NIB if necessary, and incubated at 4°C for 15min. No miracloth filtering step was performed. The cell homogenate was spun down at 4°C and 2900 x g. The resulting nuclei pellet was rinsed with 15mL NIB until the solution was clear of any photosynthetic pigments. The resulting nuclei/cellular compartment mix was used as input for the Nanobind plant nuclei big DNA kit from PacBio. Steps were followed as listed in the kit protocol except for large cell inputs, in which reagent volumes were doubled and the Proteinase K digestion step was extended to 2hrs. The isolated DNA from this protocol was processed with the Short Read Eliminator kit from PacBio to deplete DNA fragments < 25kb in length. The final, HMW DNA was used as input for nanopore library preparation.

#### Nanopore library preparation and sequencing

For all sequencing runs from axenic cultures of *E. clementina*, 1-2µg of HMW DNA was used as input to the Oxford Nanopore Technology sequencing by ligation kit (SQK-LSK112). The nanopore protocol (Version: GDE_9141_v112_revC_01Dec2021) was followed with the following minor modifications: end repair incubation was lengthened to 30min at 20°C and the adapter ligation incubation was lengthened to 60min at room temperature. Resulting libraries were loaded onto primed, high-accuracy MinION R10.4 flow cells (FLO-MIN112) at a target amount of 9 fmoles of 10kb DNA. In actuality, DNA sizes ranged within samples and between sequencing runs, but 9 fmole maintains a recommended loading amount for the flow cell at a range of fragment sizes. All sequencing of xenic cultures was performed similarly, but with previous iterations of the sequencing kit (SQK-LSK110) and the flow cell MinION R9.4.1 (FLO-MIN111). If pore occupancy dipped below roughly 1/3 of starting occupancy during the sequencing run, the run was paused, and the flow cell was washed with the Flow Cell Wash Kit (EXP-WSH004) from Nanopore according to the associated protocol. The same prepped library was then reloaded onto the flow cell and the sequencing run was restarted. Each run was left to sequence for 3-5 days, or until pore occupancy was near zero.

#### Isolation of genomic DNA for Illumina sequencing

DNA was extracted from axenic *E. clementina* cultures following the QIAGEN DNeasy Plant Pro Kit (69206) protocol. For the lysis step, cell suspension was transferred to the kit’s tissue-disrupting tubes included along with 100mg 0.5mm autoclaved glass beads added and placed in a bead-beater and shaken for one minute. 300ng of the isolated axenic *E. clementina* DNA was used as input to the NEBNext Ultra II FS DNA Library Kit for Illumina (E7805S). A fragmentation time of 16min was used for a target insert size of 200-450bp. Samples were indexed with NEBNext Multiplex Oligos for Illumina Dual Index Primers Set 1 (E7600S). DNA concentration of resulting libraries was determined with a Qubit dsDNA Quantification Assay High Sensitivity kit (Q32851). Final libraries were checked for quality and size-range using an Agilent Bioanalyzer High-Sensitivity DNA chip. The final mean insert size was 440bp, with a well-formed size distribution around the mean and minimal adapter dimers. The library was sequenced on an Illumina NextSeq 2000 P3 for 2 x 150bp reads. Raw reads were trimmed and paired with fastp (--qualified_quality_phred 20, --unqualified_percent_limit 20) for a final total of in 402 million read pairs from axenic *E. clementina*^76^.

#### RNA isolation and sequencing

To capture a wide range of transcripts, axenic cultures of *E. clementina* were exposed to different nitrogen conditions and collected at different times in the day-night cycle. Axenic cultures of *E. clementina* were seeded in 175cm^2^ sterile vented flasks at a density of 1.2 million cells per flask. For conditions of nitrogen repletion, media in the flask contained 100µM of ammonium. Cells were kept in -N or +NH_4_^+^ conditions for 72 hours and harvested two hours into the day period. Cells in nitrogen depleted conditions were additionally collected two hours into the night period. All cells were scraped from the flask, centrifuged to concentrate, resuspended in trizol, and flash frozen. Each condition was collected in triplicate for each experiment, and the whole experiment was performed twice. To lyse, the trizol suspended cells were held on ice and a sonicator probe was submerged at the center of the tube. Sonication was performed with a microtip at 50/50 on/off pulses for one minute at an intensity setting of six on a Branson 250 Sonifier (B250S). RNA was isolated using the QIAGEN RNeasy Plus Universal Mini Kit (74134) following the included protocol. 500ng of RNA per condition per replicate collected from the first experiment was used as input to the NEBNext poly(A) mRNA Magnetic Isolation module (E7490L) to enrich for mRNA and the NEBNext Ultra II Directional RNA Library Prep Kit for Illumina (E7760L) and indexes from NEBNext Multiplex Oligos for Illumina Dual Index Primers Set 1 (E7600S) were used for library preparation. 350ng of RNA per condition per replicate collected from the second experiment was used as input to the Zymo-Seq RiboFree Universal cDNA Kit (R3001) and indexed with Zymo-Seq UDI Primer Set (Indexes 1-12) (D3008). For each experiment, libraries were pooled and sequenced on an Illumina NextSeq 2000 P3 for 2 x 150bp reads.

*E. pelagica* cultures provided by courtesy of Chris Schvarcz and Kelsey McBeain proved to be unculturable long-term in lab conditions after shipment. Therefore, RNA was extracted upon receipt of overnight shipment from University of Hawaii at Manoa, HI. Otherwise, the same method of poly(A)- enrichment and Illumina sequencing was used as for *E. clementina*.

#### Data filtering and genome assembly of *E. clementina*

Initial genome size, ploidy, and repeat content estimates were made by counting k-mers in the axenic Illumina reads with jellyfish v2.2.10 (-C -m -k 35 -s 5G) and plotting with GenomeScope^77,78^. The raw fast5 sequencing files were basecalled with guppy v1.1.alpha13-0-g1ec7786. Reads were filtered based on minimum length 3kb and quality 20 with Nanofilt v2.8.0^79^. Read statistics were calculated with NanoPlot v1.30.1. Basic quality checks were performed with fastqc v0.11.9^80^. Post filtering, 19.5Gb of sequence from axenic cultures of *E. clementina* and 30.2Gb of sequencing data from xenic cultures were used for a two-step assembly process. First, axenic reads were assembled with NextDenovo v2.5.0^81^. Then, xenic nanopore sequencing data was aligned to the axenic assembly using minimap2 (-ax map-ont) v2.24-r1122 to identify probable diatom reads in the xenic data^82^. Finally, axenic and diatom-mapped xenic nanopore reads were combined and assembled with NextDenovo (using default or machine-specific options, except read_cutoff=5k, genome_size=350M). Axenic Illumina data was mapped to the assembly with BWA v0.7.17-r1188^83^. Contigs in the assembly were removed if less than 70% of the contig was covered by the axenic Illumina reads or if those reads mapped at significantly lower depth than to the rest of the contigs (< 4% of mean depth). The axenic Illumina reads were then used as input for 3 rounds of polishing with Racon v1.5.0 and one round of Polca (part of MaSuRCA v4.0.5)^84–86^. Further contamination analysis of the assembly and reads was performed with blobtools v1.1.1^87^. Organellar genomes for the diazoplast, chloroplast, and mitochondria were assembled and annotated as previously reported^28^. All contigs in the assembly were aligned to the organellar genomes and to the diazoplast genome to check for remaining organellar contaminants in the assembly. Any remaining organellar contigs contaminating the nuclear assembly were identified and removed if they aligned end-to-end to the already assembled organellar and endosymbiont genomes. Basic assembly statistics were extracted with QUAST v5.2.0^88^. Final assembly completeness and consensus quality (QV) was assessed with the k-mer spectra tool Merqury v1.3^89^. The QV of our final assembly was 38.52. BUSCO v5.3.2 in genome mode was also used to estimate completeness at the gene level^90^.

#### Repeat masking of E. clementina and E. pelagica

The final nuclear assembly of *E. clementina* and the publicly released^42^ but raw nuclear sequence of *E. pelagica* (uoEpiScrs1.2 GCA_946965045.2) were used as input to the RepeatModeler2 and RepeatMasker pipelines. To identify and classify the repeat elements for both *Epithemia* genomes, the workflow for RepeatModeler v2.0.2 with built-in LTR detection and classification was run^91^ (BuildDatabase -engine rmblast, RepeatModeler -LTRStruct, RepeatClassifier -engine rmblast). Since the repeat models for the organisms are *de novo* and repeat data for diatoms in the source databases may be limited, the repeat families classified as ‘Unknown’ were further interrogated to ensure no protein-coding genes were annotated as repeats. To do this, the Unknown repeat families were used as input to ncbi-blast+ against the NCBI non-redundant (NR) protein database^92,93^ (November 3^rd^ 2022). Approximately 8% of Unknown repeats had significant similarity to eukaryotic and Bacillariophyta proteins. Out of caution, these regions annotated as Unknown repeats with protein hits were removed from the repeat database to be kept unmasked. Finally, RepeatMasker v4.1.2-p1 (-engine rmblast, -s no_is -norna -gff -xsmall) was run on the genomes to soft-mask all repeat regions. The ParseRM tool by Aurelie Kapusta was used to extract repeat type and divergence from consensus from the raw .classified and .align output files from RepeatMasker^94^.

#### Gene annotation of E. clementina and E. pelagica

For both *E. pelagica* and *E. clementina,* the masked nuclear genome of each organism was annotated in two independent runs of BRAKER2 v2.1.6, which applied installs of GeneMark-ETP v1.0, AUGUSTUS v3.4.0, and ProtHint v2.6.0. First, the BRAKER2 pipeline was given extrinsic protein evidence as input. Protein sequences were sourced from the orthoDB v10 protozoa database which was manually edited to include diatom proteins from recent annotations. Second, the BRAKER2 pipeline was given transcriptomic evidence from the source organism^95–104^. To produce the aligned RNA-seq evidence, the RNAseq reads were quality filtered, trimmed and paired with fastp^76^ v0.22.0 (--qualified_quality_phred 20, --unqualified_percent_limit 20), and then aligned to the source genome with hisat2 v2.1.0^105^ (--rna-strandness RF). Alignment files were sorted and converted to binary alignment files with samtools v1.16.1^106^. For *E. pelagica*, a single 280 million read Illumina run from polyA-enriched RNA was used as input. For *E. clementina*, actively maintained lab cultures enabled more extensive sequencing of the transcriptome. RNA from 30 samples and five different conditions using both polyA-enrichment and rRNA-depletion methods of isolation were used as input. The outputs of these two independent protein-based and transcriptome-based annotations were merged using TSEBRA v1.0.3 into a single annotation^98^. Both the input and output general transfer format (GTF) file were fixed with the fix_gtf_ids.py script included with TSEBRA. The output GTF files were converted to multi-isoform fasta files, removing any pseudo genes or genes interrupted by stop codons using gffread v0.12.7^107^ (-J --no-pseudo -y). Completeness of the final annotation was assessed with BUSCO v5.3.2 in proteins mode. To inspect isoforms, the AGAT v1.0.0 agat_sp_keep_longest_isoform.pl tool was used^108^. Functional annotation was performed with eggNOG-mapper^109,110^. For each species, genes in shared, *Epithemia*-specific orthogroups were used for tests of enrichment against a gene universe of functionally annotated genes with p-value cutoff 0.1. clusterProfiler^111^ was used for tests of significance. Proportion of each COG term assigned within each set of functionally annotated genes was calculated and used to calculate difference in proportion.

#### Orthologue analysis

Curated species proteomes and genomes were downloaded from NCBI or associated online repositories^42,45,112–120^ (Table S1). The agat package was used to remove short isoforms (agat_sp_keep_longest_isoforms.pl). Where necessary, gene feature files were reformatted^108^ (agat_sp_manage_attributes.pl -p gene -att transcript_id). Finally, longest isoform proteomes were produced from the gene feature files and the corresponding species genome with gffread, removing genes without a complete, valid coding sequence and removing pseudo-genes^107^ (gffread -J –no-pseudo -y). The resulting proteomes were used as input for Orthologue analysis.

Orthogroups were identified with orthofinder v2.5.4 (-M msa -T iqtree) and orthogroup overlaps between species were extracted from Orthogroups_SpeciesOverlaps^121^. To quantify shared orthologues between species without biasing for total proteome size differences, the Jaccard similarity coefficient for each species pair was calculated according to the standard Jaccard index formula where A and B are the total number of self-orthologues identified for each organism and A ∩ B is the number of orthologues identified between the organisms as contained in the OrthologuesStats_one-to-one file. To identify uniquely overlapping orthogroups (e.g. orthogroups shared between two species and not by any other species), orthogroup sets from Orthogroups.GeneCount.tsv were parsed and plotted with UpSetR^122^. To quantify sequence similarity, orthologue pairs were identified by reciprocal best BLAST between organism pairs and the full-length percent amino acid identity was calculated from the BLAST outputs, similar to the method used in^123^.

#### NUDT homology search

Whole genomes of free-living cyanobacterial relatives of the endosymbiont were curated along with available whole endosymbiont genomes (Table S1). These sequences were used as query for homology searches against the nuclear genomes of *E. pelagica* and *E. clementina*. Command line BLASTN with defaults, BLASTN using the custom settings previously validated for NUMT search^92,124^ (- reward 1 -penalty -1 -gapopen 7 -gapextend 2), minimap2 (-ax asm5 and -HK19 modes), and nucmer v4.0.0rc1 were all used to perform these homology searches^82,125^. As negative controls, the reversed sequence of the *E. pelagica* mitochondria and the *E. coli* genome were used. For all cyanobacterial and endosymbiont queries, BLASTN was the most sensitive and least stringent, identifying all homology regions identified by other programs. Contiguous regions of homology < 500bp in length were not considered, though most short alignments were < 100bp. The resulting > 500bp contiguous regions of homology were considered candidate NUDTs. Seven regions in total for *E. clementina* and none in *E. pelagica.* To verify that these alignments were not a result of misassembly, long-reads from nanopore sequencing of axenic cultures were aligned (minimap -ax map-ont) and the reads spanning the border of the insertion were counted and the depth compared to that of the contig. The read depth for the contig was calculated with samtools depth (considering only primary alignments to minimize skews from repetitive regions). To check for expression within the NUDTs, RNA-seq data from both polyA enrichment and rRNA depletion experiments was mapped as previously described and normalized with deeptools v3.3.1 bamCoverage (--normalizeUsing BPM -p max -bs 100)^126^. Corresponding source regions from the endosymbiont and percent identities were pulled from the blast results. Using bedtools v2.30.0 intersect, the source regions were overlapped with endosymbiont gene regions^127^. These coordinates were then mapped back to the nuclear region. Nuclear and diazoplast sequences corresponding to these identified gene fragment containing regions were aligned using EMBOSS Needle v6.6.0.0, which calculates percent identity^128^. The truncation was calculated by dividing the length of the gene fragment by the total length of the corresponding diazoplast gene. For both the nuclear and diazoplast genomes of *E. clementina*, GC content variation was analyzed in sliding windows of 5000bp with a step size of 1000bp using bedtools makewindows and bedtools nuc. All alignments were visualized with the Integrative Genomics Viewer^129^ (IGV) and plotted with circlize^130^.

#### Identification of Horizontal Gene Transfers

Diatom proteomes (Table S1) including the *de novo* predicted *E. pelagica* and *E. clementina* were used as input to a custom HGT pipeline adapted from^56^. In brief, the program uses diamond v2.0.14 to collect homologues from the NCBI NR database for each gene in an organism^131^. To best ensure representation of genes from a diverse range of taxa, three diamond runs were performed against different subsections of NR: Bacteria, the SAR supergroup, and the remainder of the database. These results are parsed so that, where possible, the final list of homologues for each gene consists of no more than 70% of any one kingdom and does not contain any hits to self (relevant for diatom proteins already in the NCBI NR). Proteins with under 10 identifiable homologues were excluded from further analysis. Proteins were aligned with mafft v7.525 (--auto) and poorly aligned regions trimmed with trimAl v1.4.rev15 (-automated1)^132,133^. The L-INS-i method in mafft was selected for most alignments. These alignments were used as inputs for generation of phylogenetic trees. FastTree v2.1.1 was used to construct trees^134^. The topology of these trees was parsed by PhySortR v1.0.8 (min.support = 0.7, min.prop.target = 0.7, clade.exclusivity = 0.9) to identify trees in which the diatom gene of interest is more closely related to bacterial homologues than to eukaryotic ones^135^. The results were parsed using custom scripts to remove genes with fewer than five bacterial taxa in the tree. PhySortR designates genes as All Exclusive, Exclusive, Non-Exclusive, or Negative based on the tree topology. We treated All Exclusive and Exclusive results as high confidence and Non-Exclusive as low confidence. In reality, HGT candidates with Non-Exclusive tree topology are a mix of ambiguous topology as well as likely real HGTs shared between the diatoms or other eukaryotes. The Non-Exclusive HGT candidates were further filtered based on Alienness score (AI)^136^. The alienness score was calculated with both the best prokaryote Evalue and with the best prokaryote Evalue after the first group of *Bacillariophyta* results, to account for HGTs that may be shared between diatom species. HGT candidates with positive AI scores were kept for subsequent analysis. Species of origin for HGT candidates was inferred using the taxonomic breakdown of the top blast result.

#### Diazoplast Isolation

*E. clementina* cells were harvested by scraping, then washed twice in CSI-N growth medium by centrifugation at 2,000xg, and re-suspensed in spheroid body isolation buffer (50 mM HEPES pH 8.0, 330 mM D-sorbitol, 2 mM EDTA NaOH pH 8.0, 1 mM MgCl_2_). Cells were then placed in a bath sonicator for 10 minutes followed by 3 low pressure cycles (500 psi) and by 5 high pressure cycles (2,000 psi) in an EmulsiFlex-C5 Homogenizer (Avestin) or until most cells appeared lysed under a microscope. After a 1-minute spin at 100xg to pellet the unbroken cells and broken frustules, the supernatant was collected and centrifuged at 3,000xg for 5 minutes to concentrate the diazoplasts and other organelles to a volume of 3-4 mL. This fraction was then split equally, and each half was laid on a discontinuous Percoll gradient. 89% Percoll, 10% 10xPBS, and 1% 1M HEPES pH 8.0 was diluted with SIB to generate the gradient, which consisted of 2 mL 90%, 3 mL 70%, 3 mL 60%, 3 mL 50%. The gradient was centrifuged for 20 minutes at 12,000xg, 4° using a Beckman Optima L-90K ultracentrifuge with SW-41 rotor.

The boundaries between the 60% and 70% layers and the 70% and 90% layers were collected, counted, and checked for purity via light microscopy. They were then diluted 1:6 in SIB Buffer and centrifuged at 2,000xg for 2-3 minutes to collect diazoplasts, which were resuspended in 200 µL Extraction Buffer (100mM Tris-HCl, pH8.0, 2% (wt/vol) SDS, 5mM EGTA, 10mM EDTA, 1mM PMSF, 2x protease inhibitor (1 tablet each of cOmplete™ Protease Inhibitor Cocktail, catalog number 4693116001 and Pierce™ Protease Inhibitor tablet, EDTA free, catalog number A32965)). During optimization, enrichment was assessed by Western blot for NifDK on both the diazoplast and whole cell extracts.

#### Protein Extraction, Preparation, and LCMS/MS

We generated whole cell lysate by homogenizing with a bead beater at 3000 strokes per minute for 3 minutes with 1 mm glass beads (BioSpec Products catalog number 11079110) or until most cells appeared lysed under a microscope. Diazoplasts were lysed similarly using 0.5 mm beads (BioSpec Products catalog number 11079105). Beads were pelleted at 100xg for 1 minute and the supernatant was removed; the beads were washed twice with 50 µL extraction buffer each by vortexing and spinning. These fractions were then added to the supernatant for a total of 300 µL, followed by an equal volume of cold Tris-buffered phenol (pH 7.5-7.9). This solution was vortexed for 1 minute, centrifuged at 18,000 x g for 15 minutes at 4° C. The upper phase was discarded, then extracted with an equal volume of cold 50mM Tris-HCl, pH8.0. The phenol phase was extracted with Tris-HCl a total of three times, followed by addition of 0.1 M ammonium acetate in methanol and overnight incubation at -80° C. Samples were then transferred to new tubes and centrifuged at 18,000 x g for 20 minutes at 4° C. The supernatant was discarded and the pellet was washed once in 0.1 M ammonium acetate in methanol and twice in 1 mL cold methanol by centrifugation for 5 minutes at 18,000 x g at 4° C, followed by a final short spin and removal of trace methanol. The pellet was then resuspended in 150 µL resuspension buffer (6M Guandine-HCl in 25mM NH_4_HCO_3_ pH8.0). Each sample was then reduced with TCEP at a final concentration of 2 µM (Thermo Scientific catalog number 20490) for 1 hour at 56° C, alkylated with iodoacetamide (Thermo Scientific catalog number 90034) at a final concentration of 10 mM for 1 hour at ambient temperature, and then diluted with 3 volumes of 25mM NH4HCO3. Sequencing grade modified trypsin (Promega catalog number V5111) was added at a ratio of 1:50 followed by overnight incubation at 37° C, then repeated the next morning, followed by quenching the reaction by adding formic acid to a final concentration of 1%. Each sample was then loaded onto a C18 cartridge (Sep-pak waters catalog number WAT054960), activated with 80% acetonitrile and 0.1% formic acid. The flow-through was loaded a total of three times, followed by five washes with 1 mL 0.1% formic acid. The samples were then eluted with 200 µL of 80% acetonitrile 1% formic acid and the flow-through re-loaded a total of three times.

Peptide concentration was determined using Pierce™ Quantitative Colorimetric Peptide Assay (Thermo Fisher catalog number 23275). 1 µg of peptides from each sample was loaded on either on a Q-Exactive HF hybrid quadrupole-Orbitrap mass spectrometer (Thermo Fisher) (1 replicate) or an Eclipse Tribrid mass spectrometer (Thermo Fisher) (2 replicates), equipped with an Easy LC 1200 UPLC liquid chromatography system (Thermo Fisher). Peptides were first trapped using a trapping column (Acclaim PepMap 100 C18 HPLC, 75 μm particle size, 2 cm bed length), then separated using analytical column AUR3-25075C18, 25CM Aurora Series Emitter Column (25 cm x 75 µm, 1.7µm C18) (IonOpticks). The flow rate was 300 nL/min, and a 120-min gradient was used. Peptides were eluted by a gradient from 3 to 28 % solvent B (80 % acetonitrile, 0.1 % formic acid) over 106 min and from 28 to 44 % solvent B over 15 min, followed by a short wash (9 min) at 90 % solvent B. The Q-Exactive HF hybrid quadrupole-Orbitrap mass spectrometer was configured as follows: Precursor scan was from mass-to-charge ratio (m/z) 375 to 1600 (resolution 120,000; AGC 3.0E6, maximum injection time 100ms) and top 20 most intense multiply charged precursors were selected for fragmentation (resolution 15,000, AGC 5E4, maximum injection time 60ms, isolation window 1.0 m/z, minimum AGC target 1.2e3, intensity threshold 2.0 e4, include charge state =2-8). Peptides were fragmented with higher-energy collision dissociation (HCD) with normalized collision energy (NCE) 27. Dynamic exclusion was enabled for 24s. The Orbitrap Eclipse Tribrid mass spectrometer was configured as follows: Precursor scan was from mass-to-charge ratio (m/z) 375 to 1600 (resolution 120,000; AGC 200,000, maximum injection time 50ms, Normalized AGC target 50%, RF lens(%) 30) and the most intense multiply charged precursors were selected for fragmentation (resolution 15,000, AGC 5E4, maximum injection time 22ms, isolation window 1.4 m/z, normalized AGC target 100%, include charge state=2-8, cycle time 3 s). Peptides were fragmented with higher-energy collision dissociation (HCD) with normalized collision energy (NCE) 27. Dynamic exclusion was enabled for 30s.

#### Proteomics Data Analysis

Maxquant version 2.5.0 was used for proteomics database searches, using default parameters with the following changes: label-free and iBAQ quantification, matched between runs were enabled^137^. For identifications, peptides were searched against the *Epithemia clementina* reference host and diazoplast proteomes. The proteingroups.txt file output from MaxQuant was analyzed using Perseus version 2.0.10.0^138^. iBAQ values were imported and filtered to remove potential contaminants, reverse hits, and those only identified by site. Only proteins identified by two more unique peptides and with a minimum of 5% sequence coverage were included in further analysis. The iBAQ values were then log(2) transformed for normality, proteins with two or more non-valid values were removed, and missing values were imputed from a downshifted normal distribution of the total matrix (width 0.3 standard deviations, down shift 1.8 standard deviations). A two-sided students T-test using the significance analysis of microarrays method (s0=0.1, false discovery rate 0.05, 250 randomizations) was used to determine the enrichment of host-encoded proteins in the diazoplast.

#### Immunoblot

Whole cell and isolated diazoplast lysates were prepared as described above. Protein concentration was determined using Pierce™ BCA Protein Assay Kit (Thermo Fisher catalog number 23227). 0.5 µg of protein from each sample was diluted in lithium dodecyl sulfate buffer with 100 mM DTT and loaded onto a NuPage™ Bis-Tris gels 4-12% acrylamide (Thermo Fisher catalog number NP0321BOX) in MES Buffer, and separated by electrophoresis, using Precision Plus Protein™ All Blue Standards (BioRad catalog number 1610373). Proteins were then transferred into a nitrocellulose membrane using Bio-rad Transblot Turbo, followed by blocking in LiCOR blocking buffer 1 hour at room-temperature. The membrane was then incubated for two hours at room temperature with primary antibodies anti-NifDK (polyclonal goat at 1:5000 dilution, kindly provided by Dr. Dennis Dean from Virginia Tech, US) to detect nitrogenase and anti-PsbA (1:10,000 dilution rabbit from AgriSera AB, Vanas, Sweden) as an internal loading control. Antibodies were diluted in a solution of 50% TBST and 50% LiCOR blocking buffer. The membrane was then washed three times with TBST and incubated with LiCOR secondary antibodies (IRDye 800CW) for 1 hour (goat α-rabbit for PsbA and donkey α-goat for NifDK). After two rinses with TBST and one with PBS, the blot was imaged using an infra-red LiCOR imager. Intensity of the signal was quantified using Image Studio Lite software v5.2.

### QUANTIFICATION AND STATISTICAL ANALYSIS

In all cases, statistical analyses were performed using pre-existing software. The specific statistical tests, settings, and cutoffs used are listed with their accompanying analysis in the preceding method details.

