## Supplemental Data S1 A-G for "Genomes of nitrogen-fixing eukaryotes reveal a non-canonical model of organellogenesis"

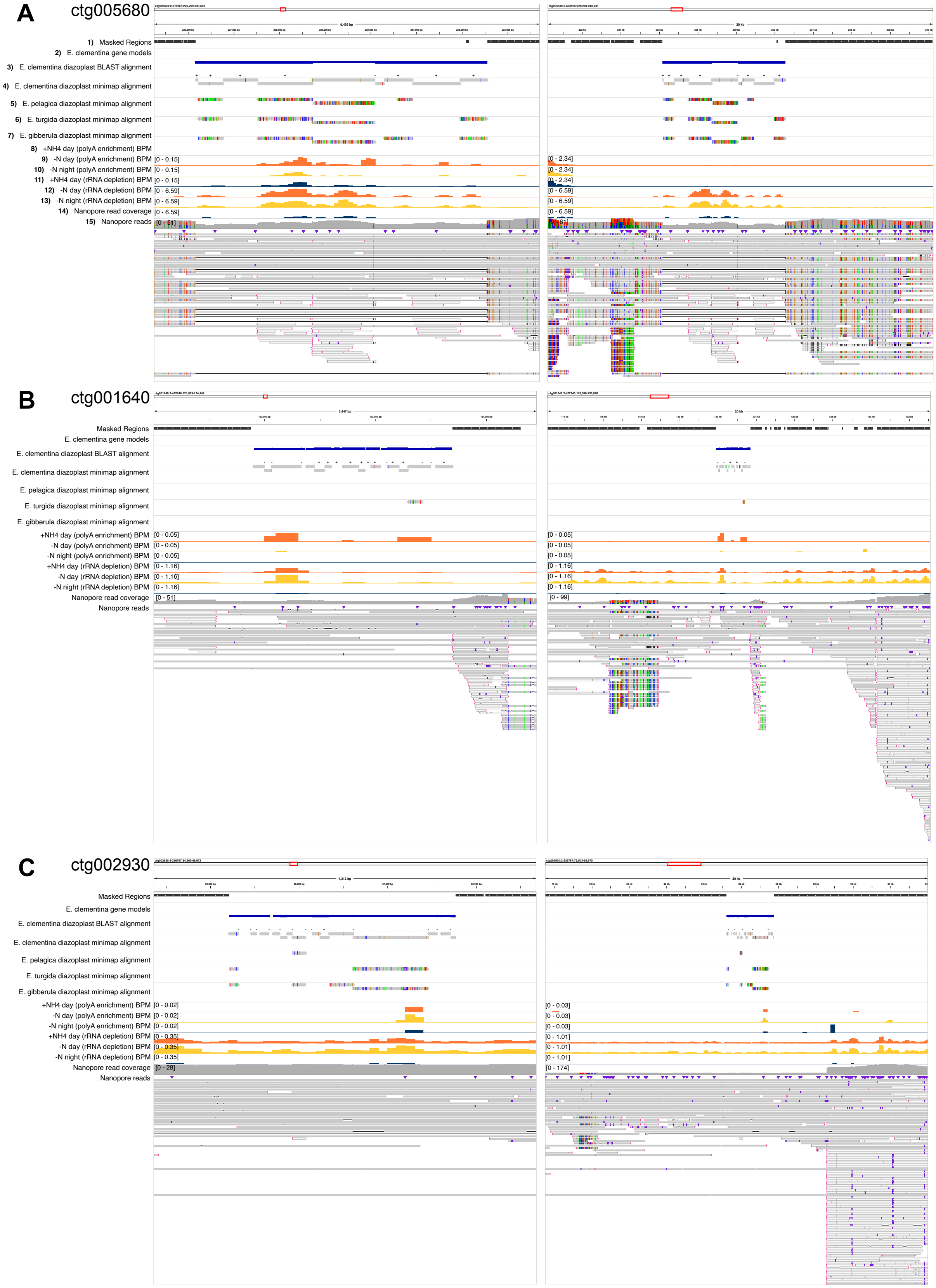

A
ctg005680
1) Masked Regions
2) E. clementina gene models
3) E. clementina diazoplast BLAST alignment
4) E. clementina diazoplast minimap alignment
5) E. pelagica diazoplast minimap alignment
6) E. turgida diazoplast minimap alignment
7) E. gibberula diazoplast minimap alignment
8) +NH4 day (polyA enrichment) BPM
9) -N day (polyA enrichment) BPM
10) -N night (polyA enrichment) BPM
11) +NH4 day (rRNA depletion) BPM
12) -N day (rRNA depletion) BPM
13) -N night (rRNA depletion) BPM
14) Nanopore read coverage
15) Nanopore reads
[0 - 0.15]
[0 - 6.59]
[0 - 2.34]
B
ctg001640
Masked Regions
E. clementina gene models
E. clementina diazoplast BLAST alignment
E. clementina diazoplast minimap alignment
E. pelagica diazoplast minimap alignment
E. turgida diazoplast minimap alignment
E. gibberula diazoplast minimap alignment
+NH4 day (polyA enrichment) BPM
-N day (polyA enrichment) BPM
-N night (polyA enrichment) BPM
+NH4 day (rRNA depletion) BPM
-N day (rRNA depletion) BPM
-N night (rRNA depletion) BPM
Nanopore read coverage
Nanopore reads
[0 - 0.05]
[0 - 1.16]
[0 - 51]
[0 - 99]
C
ctg002930
Masked Regions
E. clementina gene models
E. clementina diazoplast BLAST alignment
E. clementina diazoplast minimap alignment
E. pelagica diazoplast minimap alignment
E. turgida diazoplast minimap alignment
E. gibberula diazoplast minimap alignment
+NH4 day (polyA enrichment) BPM
-N day (polyA enrichment) BPM
-N night (polyA enrichment) BPM
+NH4 day (rRNA depletion) BPM
-N day (rRNA depletion) BPM
-N night (rRNA depletion) BPM
Nanopore read coverage
Nanopore reads
[0 - 0.02]
[0 - 0.35]
[0 - 28]
[0 - 0.03]
[0 - 1.01]
[0 - 174]

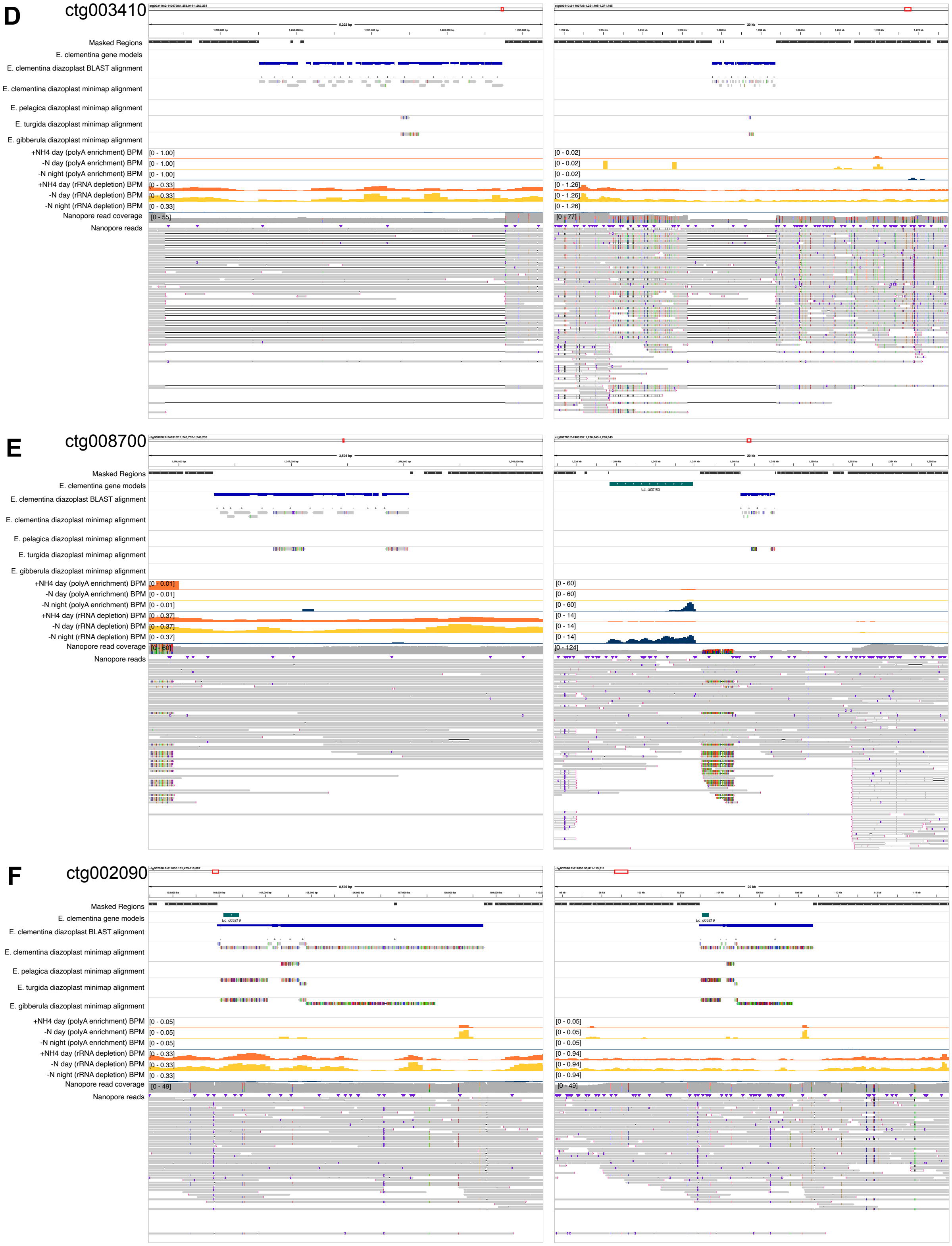
D
ctg003410
Masked Regions
E. clementina gene models
E. clementina diazoplast BLAST alignment
E. clementina diazoplast minimap alignment
E. pelagica diazoplast minimap alignment
E. turgida diazoplast minimap alignment
E. gibberula diazoplast minimap alignment
+NH4 day (polyA enrichment) BPM
-N day (polyA enrichment) BPM
-N night (polyA enrichment) BPM
+NH4 day (rRNA depletion) BPM
-N day (rRNA depletion) BPM
-N night (rRNA depletion) BPM
Nanopore read coverage
Nanopore reads
[0 - 1.00]
[0 - 1.00]
[0 - 1.00]
[0 - 0.33]
[0 - 0.33]
[0 - 0.33]
[0 - 55]
[0 - 0.02]
[0 - 0.02]
[0 - 0.02]
[0 - 1.26]
[0 - 1.26]
[0 - 1.26]
[0 - 77]
E
ctg008700
Masked Regions
E. clementina gene models
E. clementina diazoplast BLAST alignment
E. clementina diazoplast minimap alignment
E. pelagica diazoplast minimap alignment
E. turgida diazoplast minimap alignment
E. gibberula diazoplast minimap alignment
+NH4 day (polyA enrichment) BPM
-N day (polyA enrichment) BPM
-N night (polyA enrichment) BPM
+NH4 day (rRNA depletion) BPM
-N day (rRNA depletion) BPM
-N night (rRNA depletion) BPM
Nanopore read coverage
Nanopore reads
[0 - 0.01]
[0 - 0.01]
[0 - 0.01]
[0 - 0.37]
[0 - 0.37]
[0 - 0.37]
[0 - 60]
Ec_g22162
[0 - 60]
[0 - 60]
[0 - 60]
[0 - 14]
[0 - 14]
[0 - 14]
[0 - 124]
F
ctg002090
Masked Regions
E. clementina gene models
E. clementina diazoplast BLAST alignment
E. clementina diazoplast minimap alignment
E. pelagica diazoplast minimap alignment
E. turgida diazoplast minimap alignment
E. gibberula diazoplast minimap alignment
+NH4 day (polyA enrichment) BPM
-N day (polyA enrichment) BPM
-N night (polyA enrichment) BPM
+NH4 day (rRNA depletion) BPM
-N day (rRNA depletion) BPM
-N night (rRNA depletion) BPM
Nanopore read coverage
Nanopore reads
Ec_g05219
[0 - 0.05]
[0 - 0.05]
[0 - 0.05]
[0 - 0.33]
[0 - 0.33]
[0 - 0.33]
[0 - 49]
Ec_g05219
[0 - 0.05]
[0 - 0.05]
[0 - 0.05]
[0 - 0.94]
[0 - 0.94]
[0 - 0.94]
[0 - 49]

**G** ctg003780

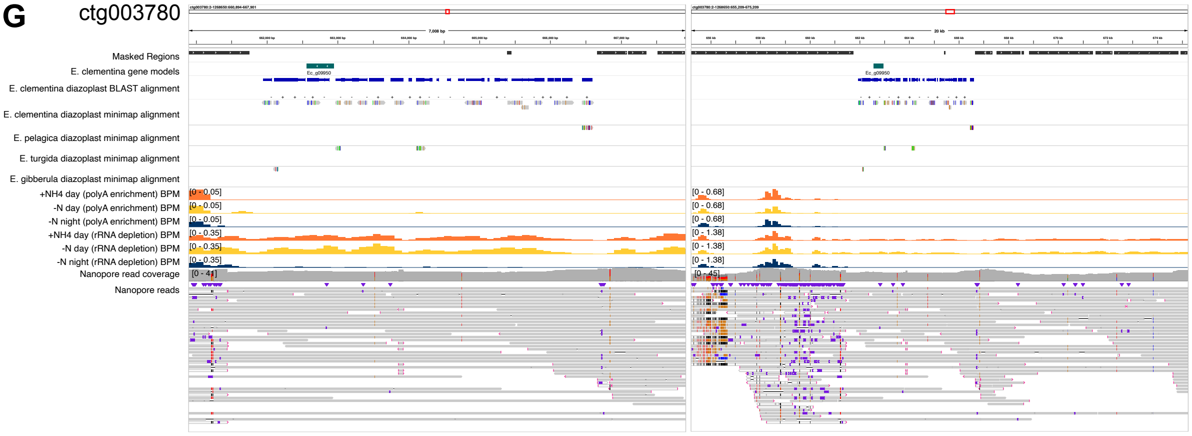


**Data S1: Detailed genome tracks across NUDT regions**

(A-G) For all NUDTs, full context genome tracks from the Integrated Genomics Viewer zoomed in to the NUDT region (left) or zoomed out to a 20kb surrounding region (right). Tracks from top to bottom are:

1) Region file of masked repeat regions;
2) Feature file of E. clementina gene models;
3) Region file of E. clementina diazoplast homology found by BLAST, demarcates the NUDT;

4-7) Alignment files of homology found by minimap2 when aligning 4) E. clementina diazoplast, 5) E. pelagica diazoplast, 6) E. turgida diazoplast, and 7) E. gibberula diazoplast to the E. clementina nuclear genome;

8-10) Normalized expression data in BPM of RNA seq from combined replicates of poly-adenylated transcript enriched RNA collected from three treatment conditions.

11-13) Normalized expression data in BPM of RNA seq from combined replicates across of ribosomal RNA depleted RNA collected from three treatment conditions.

14) Read pileup of axenic nanopore reads. Colored bars at certain sites indicate proportion of SNVs across the reads deviating from the haplotype reference assembly often resulting from a heterozygous site but sometimes from reads accumulating at undiscernible copies of repeat elements.

15) Alignment file of axenic, nanopore long reads aligned to the reference E. clementina genome. An aligned read identical to the reference sequence is rendered as a single plain grey bar. Colors at sites along the read denote SNVs from the reference assembly. Small indels are denoted by small purple bars. A thin black bar within a read represents a region not present in the read that is present in the haplo-assembly (i.e. larger indels). Very light grey bars are secondary alignments, which accumulate at repeat elements.
